# Distinct mechanisms underlie H_2_O_2_ sensing in *C. elegans* head and tail

**DOI:** 10.1101/2021.07.26.451501

**Authors:** Sophie Quintin, Théo Aspert, Tao Ye, Gilles Charvin

## Abstract

Environmental oxidative stress threatens cellular integrity and should therefore be avoided by living organisms. Yet, relatively little is known about environmental oxidative stress perception. Here, using microfluidics, we showed that like I2 pharyngeal neurons, the tail phasmid PHA neurons function as oxidative stress sensing neurons in *C. elegans*, but display different responses to H_2_O_2_ and light. We uncovered that different but related receptors, GUR-3 and LITE-1, mediate H_2_O_2_ signaling in I2 and PHA neurons. Still, the peroxiredoxin PRDX-2 is essential for both, and might promote H_2_O_2_-mediated receptor activation. Our work demonstrates that *C. elegans* can sense a broad range of oxidative stressors using partially distinct H_2_O_2_ signaling pathways in head and tail sensillae, and paves the way for further understanding of how the integration of these inputs translates into the appropriate behavior.

**Graphical abstract:** 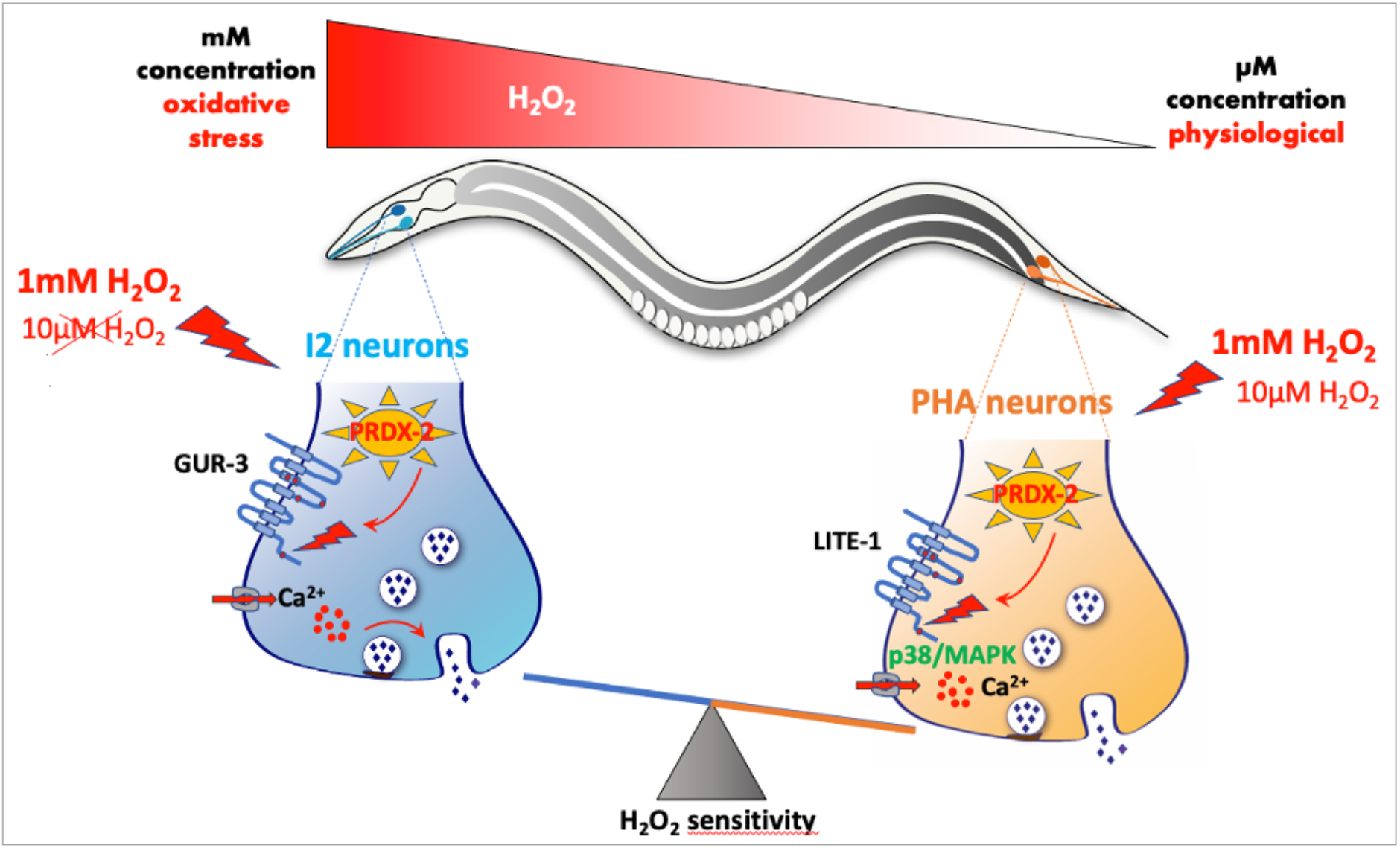

## Introduction

Reactive oxygen species (ROS) are well-known to exert a dual effect, promoting aging and pathological conditions on the one hand and increasing organism resistance and longevity on the other hand [1]. Neurons are easily exposed to ROS, and many neurodegenerative diseases have been associated with oxidative stress [2]. However, exposure of *C. elegans* nematodes to a mild oxidative stress has been reported to be beneficial for neuron sensory function: micromolar doses of the ROS-inducing agent paraquat or of hydrogen peroxide (H_2_O_2_) improve the sensitivity in ASH polymodal neurons [3,4], whereas millimolar doses of H_2_O_2_ reduce neuron response [3], likely inducing oxidative stress. Therefore, it is essential for nematodes to detect a broad range of H_2_O_2_ concentrations to preserve their cellular integrity.

Although the cellular response to oxidative stress has been extensively characterized (reviewed in [5]), little is known on how oxidants are actually perceived and which underlying molecular pathways are involved. A recent study has revealed the behavioral mechanisms which allow *C. elegans* to find a niche providing both food and protection from H_2_O_2_ [6]. In addition, light and H_2_O_2_ sensing were shown to be tightly connected in yeast [7] and in nematodes [8]. While it has been demonstrated that light is converted into an H_2_O_2_ signal in yeast [7], this question remains unanswered in nematodes. Notably, *C. elegans* can detect both H_2_O_2_ and light via the I2 pharyngeal neurons and responds to these stimuli by inhibition of feeding or by an avoidance behavior [8]. Initially described as interneurons, the I2 neurons proved to be primary sensory neurons [9] that are highly specialized in oxidative stress sensing [8], but were also shown to respond moderatly to salt or odor [10]. However, although detection of a large spectrum of H_2_O_2_ is critical for nematodes, the range of H_2_O_2_ concentrations detected by I2 neurons has not been investigated, and the molecular mechanisms involved in H_2_O_2_ signaling are not well defined.

In addition, the nematode also possesses tail sensory neurons specialized in chemo-repulsion, called PHA/PHB (or phasmids, reviewed in [11]). In contrast to I2 neurons, these tail neurons can respond to many noxious stimuli [12] and trigger avoidance [13], but whether PHA neurons can sense H_2_O_2_ or light remains an open question.

H_2_O_2_ sensing in I2 neurons requires the function of the peroxiredoxin PRDX-2 [8], a highly conserved antioxidant enzyme whose role remains unclear. Peroxiredoxins (Prxs) belong to a family of thiol peroxidases which can reduce H_2_O_2_ in cells following the oxidation of one or two cysteines in their catalytic domain, and they are highly abundant from yeast [14,15] to human cells, in which they represent approximately 1% of the total dry cellular mass [16,17]. Oxidised Prxs are recycled in the reduced, active form, by thioredoxin [18]. At high H_2_O_2_ concentrations, Prxs become hyperoxidized, a form that has been shown to function as a molecular chaperone [19,20]. Similarly, thioredoxins have been shown to have redox-independent function [21,22]. As pivotal antioxidants, Prxs dysfunction has been associated with several pathologies [23], including cancer [24]. In budding yeast, the peroxiredoxin Tsa1 has a major role in maintaining the redox balance, and is massively induced upon oxidative stress as a direct target of the H_2_O_2_-sensing transcription factor Yap1 [25,26]. In *C. elegans*, among three genes encoding peroxiredoxins, PRDX-2 is the only one whose depletion induces a phenotype and is considered as the major peroxiredoxin [27,28]. PRDX-2 is expressed in many cell types including neurons, gut [27–29], muscle and epithelial cells [8]. Although a global induction of *prdx-2* mRNA expression has been reported upon treatment with the strong tBOOH oxidant [28], cells in which PRDX-2 expression is induced remain to be identified. Likewise, the question of a tissue-specific regulation of PRDX-2 by SKN-1, the closest ortholog of Yap1, has not been addressed.

Importantly, beyond their peroxidase activity, Prxs have long been proposed to act as intracellular H_2_O_2_ sensors, which influence cellular signaling [30–32]. For example, the p38/MAPK signaling pathway, which controls adaptive mechanisms and/or cell fate decisions, is activated by H_2_O_2_ through Prxs in an evolutionary conserved manner [33,34]. Like in mammalian or in drosophila cells, several studies in *C. elegans* indicate that PRDX-2 would relay H_2_O_2_ signaling, activating the downstream p38/PMK-1 pathway [3,35,36]. Notably, low doses of H_2_O_2_ potentiate the sensory response of ASH sensory neurons to glycerol through activation of the PRDX-2/p38/PMK-1 cascade [3]. Yet, whether this cascade is at play in I2 neurons to influence H_2_O_2_ perception has not been analyzed.

Here, we undertook a subcellular analysis of the peroxiredoxin PRDX-2 in *C. elegans*, focusing on its requirement in neuronal H_2_O_2_ sensing. Using a CRISPR knock-in line, we showed that PRDX-2 is present in many cells, among which several pairs of neurons: I2s in the head, PHAs in the tail and CANs in the body. Interestingly, upon an H_2_O_2_ challenge, an upregulation of PRDX-2 is observed only in the anterior gut and in the excretory pore, but not in neurons, suggesting that PRDX-2 might fulfill different functions, depending on the cell type where it is expressed. Using a microfluidic-based approach and real-time calcium imaging, we show that PHA neurons also respond to H_2_O_2_, with an even higher sensitivity than I2 neurons. Although H_2_O_2_ perception depends on *prdx-2* function in both pairs of neurons, we uncovered that it relies on different gustatory receptors and downstream transducers: while dispensable in I2 neurons, the p38/MAPK kinase pathway contributes to the hypersensitivity of PHA neurons to H_2_O_2_. Interestingly, we uncovered that H_2_O_2_ sensing requires the same receptors as light sensing, and that PHA neurons respond to light — establishing a parallel between PHA and photosensory ASH neurons. Based on our work and on previous studies, we propose a molecular model of how H_2_O_2_ could trigger neuronal activation in I2 and PHA through a peroxiredoxin-mediated redox relay. Taken together, our data suggest that *C. elegans* can sense a broad range of oxidative stress using partially distinct H_2_O_2_ signaling pathways acting in head and tail sensillae.

## Results

### Expression pattern of PRDX-2::GFP and its variation upon H_2_O_2_ treatment

In budding yeast, the peroxiredoxin Tsa1 is massively induced upon H_2_O_2_ treatment [25,26]. To gain insight into the tissue-specific expression of PRDX-2 upon oxidative stress, we first used the PRDX-2 reporter line generated by [29]. However, the level of expression of this strain varies a lot, and transgenics show many fluorescent aggregates ([29], S1 Fig), likely associated with transgene overexpression. Consistently, the strain displays a much stronger resistance to oxidative stress than wild-type animals (S1 Fig), suggesting that overexpressed PRDX-2 construct induces a higher H_2_O_2_ scavengeing capacity in transgenics. Furthermore, it was impossible to identify PRDX-2-expressing cells in this strain, preventing the use of this strain in our study. For these reasons, we created a GFP knock-in line of PRDX-2, using the CRISPR-Cas9 technique [37]. A C-terminus GFP fusion targeting all PRDX-2 isoforms, comprising a linker, was engineered and inserted at the *prdx-2* locus (Fig 1A). Three independent knock-in lines were obtained, sharing an identical expression pattern (Fig 1B, S1 Fig).

**Figure 1.**
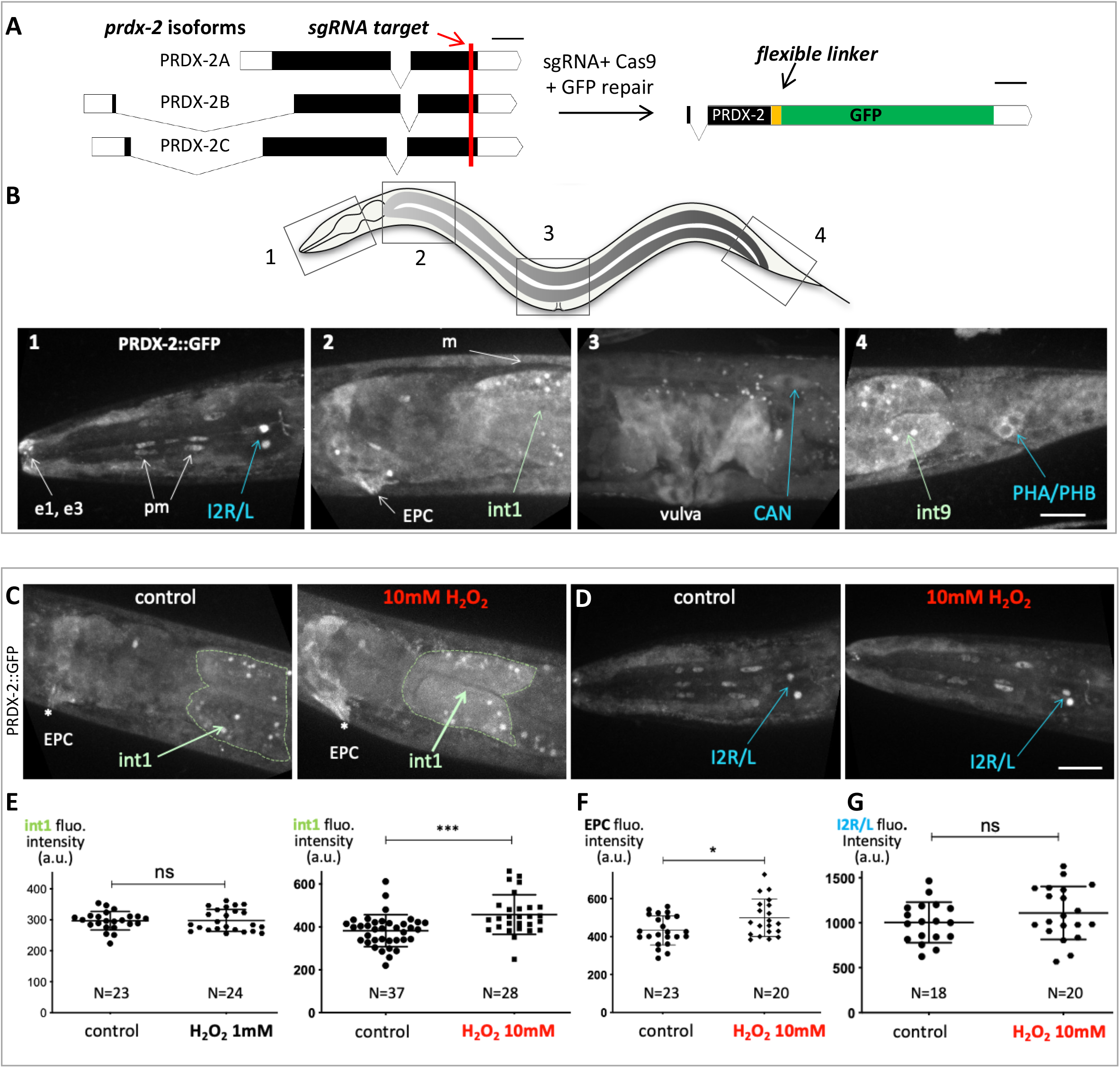
PRDX-2::GFP knock-in line expression pattern and its variation upon H_2_O_2_ treatment. (A) Sketch depicting the PRDX-2::GFP knock-in strategy using CRISPR Cas9-mediated genome editing. The sgRNA target sequence (shown in red) was chosen upstream PRDX-2 STOP codon, to tag all *prdx-2* isoforms. Black boxes indicate exons, white boxes unstranslated regions. Bar, 100 bases pairs. The resulting last exon of the PRDX-2::GFP fusion protein is shown (right); note the insertion of a flexible linker between PRDX-2 and the GFP, to promote correct folding. After injection, three independent knock-in lines were recovered, sharing the same expression pattern. (B) Spinning-disc confocal projections of a representative PRDX-2::GFP knock-in animal, in 4 body regions (boxed in the worm sketch on top). PRDX-2::GFP expression is observed in the tip of the nose (e1, e3 epithelial cells), in pharyngeal muscle cells (pm), in body wall muscles (m), in the excretory pore cell (EPC), in proximal and distal gut cells (int1 and int9), and in several neuron pairs (I2, CAN, PHA/PHB). Note the different level of PRDX-2::GFP in I2 left and right neurons in panel B1. (C-G) An acute oxidative stress triggers an upregulation of PRDX-2 in the anterior gut, but not in neurons. (C-D) Spinning-disc confocal projections of control or H_2_O_2_-treated animals in the anterior gut (C) or in the head (D). (E-G) Quantification of fluorescence intensity in controls and in H_2_O_2_-treated animals in the int1 cell (E), in the EPC (F), and I2 neurons (G). Bars indicate mean and SD. ns, not significant, p>0.05; *p<0.05; ***p<0.001 (t test and Mann Whitney test). Scale bar, 20μm.

In the PRDX-2::GFP knock-in line, we detected a broad expression of PRDX-2 in various cell types, including the proximal and distal gut, muscles (body wall, vulval and pharyngeal), epithelial cells (vulva, nose tip, hypodermis) and I2 neurons (Fig 1B), as previously reported [8,27,28]. In addition, we observed for the first time PRDX-2 expression in the excretory pore cell (EPC), and in two other pairs of neurons; the tail phasmids (PHA/PHB, whose identity was ascertained by DiI staing, S1 Fig) and the excretory canal-associated neurons (CANs), located close to the vulva (Fig 1B). Consistent with this, a high number of *prdx-2* transcripts was detected in CANs and in the EPC [38]. Therefore, we hypothesized that the knock-in line faithfully reflects the endogenous PRDX-2 expression, and characterized it further.

### PRDX-2::GFP is induced in the anterior gut upon H_2_O_2_ treatment, but not in I2 neurons

We noticed that many PRDX-2-expressing cells are directly in contact with the environment, such as the EPC, the tip of the nose, the vulva, and neurons, which all possess terminations exposed outside. This pattern is strikingly reminiscent of that detected in animals carrying the *HyPer* H_2_O_2_ biosensor after H_2_O_2_ exposure [39]. Thus, PRDX-2 is expressed in cells in which environmental H_2_O_2_ penetrates more easily, suggesting a protective role of PRDX-2 in these cells as a peroxidase. Therefore, in the following, we wondered whether all PRDX-2-expressing cells respond similarly to an H_2_O_2_-induced oxidative stress.

To determine whether PRDX-2 expression changes upon oxidative stress, we exposed animals to different doses of H_2_O_2_, and PRDX-2::GFP expression was quantified in different cells after spinning-disc confocal acquisitions. We selected H_2_O_2_ concentrations inducing different physiological responses [8]. A higher PRDX-2 expression was detected in the anterior gut two hours after a 10mM H_2_O_2_ treatment, but not after a 1mM H_2_O_2_ treatment (Fig 1C,E). The fluorescence was specific to PRDX-2::GFP as H_2_O_2_-treated controls did not show a higher gut autofluorescence (S2 Fig). Similarly, PRDX-2 expression was only induced in the EPC at 10mM H_2_O_2_ (Fig 1C,F, S2 Fig). Thus, our data indicate a dose-dependent PRDX-2 induction—as it occurs at 10mM but not at 1mM. The origin of this difference could arise from the animal behavioral response: at 1mM H_2_O_2_ the pharyngeal pumping is strongly inhibited to prevent ingestion [8], thereby exposure of gut to H_2_O_2_. This could explain both the absence of PRDX-2 induction in the gut we report, and the unchanged level of *prdx-2* mRNA reported [28], after a 1mM H_2_O_2_ treatment. In contrast, at 10mM H_2_O_2_, the nematode should retract back (avoidance response, [8]); but here, as trapped animals cannot escape, they likely swallow a certain amount of H_2_O_2_, exposing the foregut to a severe oxidative stress which could trigger PRDX-2 induction. A 10mM H_2_O_2_ treatment of 30min has been shown to induce hyperoxidation of over 50% total PRDX-2 in wild-type lysed worms [40] and this form persists after a 4h recovery period [27]. Although the induction we observed is only based on expression level and not on protein activity, we suggest that PRDX-2 could still scavenge H_2_O_2_ in the EPC and in the foregut, consistent with the reported protective role of intestinal PRDX-2 [27].

Among PRDX-2-expressing neurons, the I2 pair shows the highest expression and possesses terminations exposed to the outside [9]. Therefore, we focused on I2 neurons to investigate whether the neuronal level of PRDX-2 is affected upon H_2_O_2_ treatment. However, at both concentrations tested (1 and 10mM), the level of PRDX-2 in I2 neurons constantly remained unchanged after H_2_O_2_ treatment (Fig 1D,G, S2 Fig).

Thus, we observed an upregulation of PRDX-2::GFP in the anterior gut and in the EPC following an acute oxidative stress, but not in neurons. As Prxs belong to a homeostatic system, an upregulation was shown to be associated with a detoxification function in yeast [26]. By analogy, PRDX-2 might fulfill a peroxidase function in the foregut and in the EPC to protect the animal against environmental aggressions. In contrast, in I2 neurons, PRDX-2 would instead act in H_2_O_2_ sensing and/or signaling, as proposed [8,28]. We conclude that the responses of PRDX-2 to oxidative stress are cellular context-dependent.

### SKN-1 controls expression of PRDX-2 in the gut, but not in neurons

This prompted us to test whether PRDX-2 expression could rely on different subcellular regulations. By analogy with yeast, we wondered whether PRDX-2 would be controlled by the Yap1 nematode orthologue SKN-1 [5] in cells where PRDX-2 is induced upon oxidative stress, and asked whether such regulation occurs in other cells. We sought to test this hypothesis by inactivating the function of SKN-1 in the PRDX-2::GFP knock-in line. We first used the *skn-1(zj15)* allele, which specifically inactivates gut-specific isoforms, leaving neuronal isoforms unaffected [41]. In this mutant, the basal expression level of PRDX-2 is reduced in the foregut, but an induction persists after a 10mM H_2_O_2_ treatment (Fig 2C), suggesting that the neuronal isoforms could mediate this effect. Consistent with this, the RNAi-mediated knock-down of all *skn-1* isoforms also triggered a reduction of PRDX-2::GFP signal in the anterior gut, and an absence of induction upon H_2_O_2_ treatment (Fig 2A,B,D). This result indicates that SKN-1 activity is essential to regulate PRDX-2 expression in the anterior gut, both at the basal level and under oxidative stress. In agreement with this, SKN-1 was found to bind to the *prdx-2* promoter in chromatin immunoprecipitation experiments coupled with high-throughput DNA sequencing (ChIP-seq) [42] (S3 Fig).

**Figure 2.**
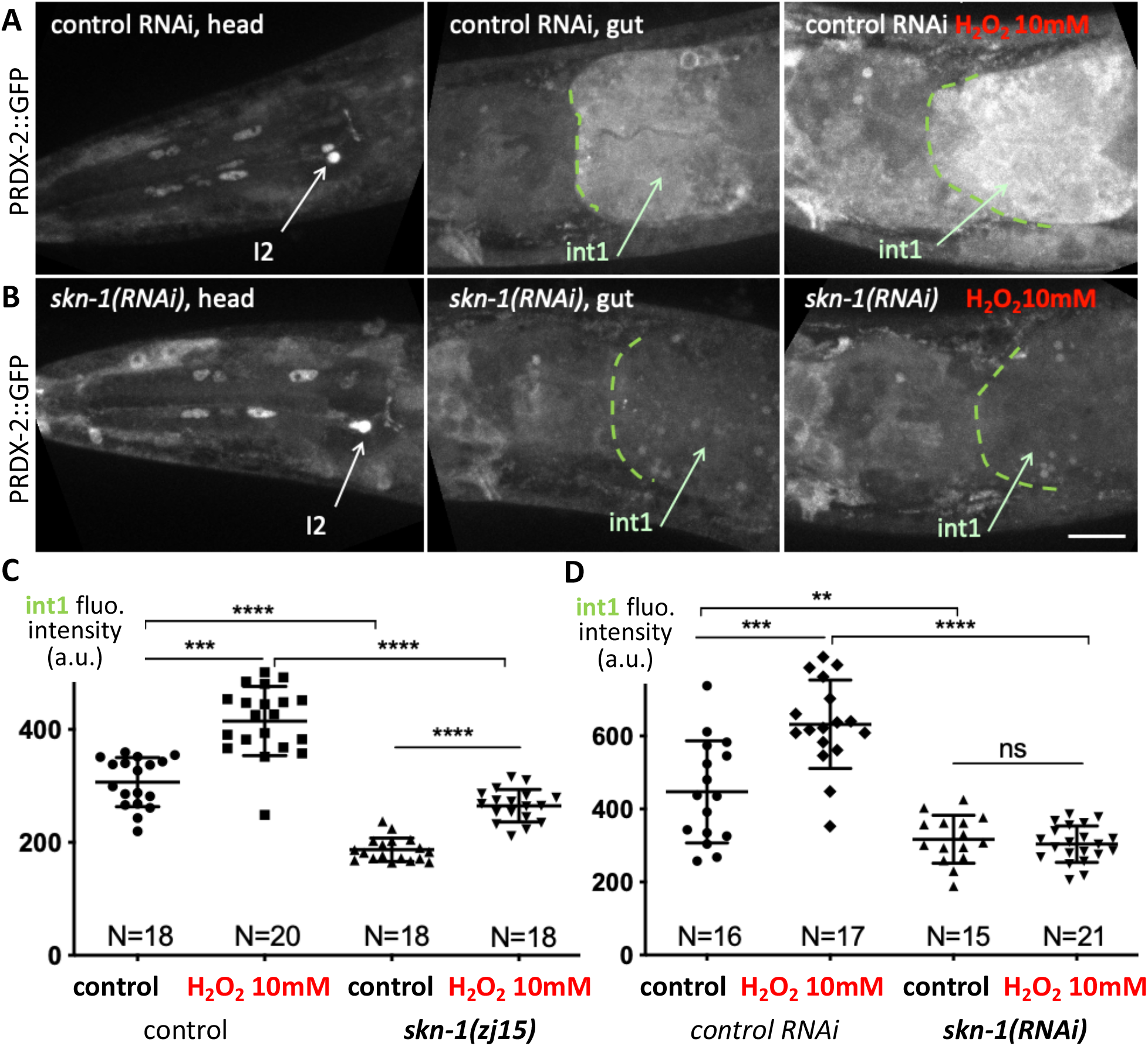
SKN-1 function is required for PRDX-2 expression in the gut. (A-B) Spinning-disc confocal projections of head and anterior gut of PRDX-2::GFP knock-in animals, in control RNAi (A) and in *skn-1(RNAi)* (B) animals. The right panel shows the foregut of a 10mM H_2_O_2_-treated animal in both genotypes. Note the low level of intl PRDX-2::GFP fluorescence in *skn-1(RNAi)* animal. (C-D) Quantification of the int1 cell fluorescence intensity in control and after a 10mM H_2_O_2_-treatment, in *skn-1(zj15)* mutants (C) and in *skn-1(RNAi)* animals (D). Bars indicate mean and SD. ns, not significant, p>0.05; **p<0.01; ***p<0.001; ***p<0.0001 (t test, ANOVA). Scale bar, 20μm.

In contrast, in I2 neurons, no change in the PRDX-2 level was detected at the basal level or after H_2_O_2_ treatment compared to controls, neither in *skn-1(zj15)* mutants nor in *skn-1(RNAi)* animals (S2 Fig). With the limitation that RNAi efficiency may be lower in neurons [43], we suggest that additional transcription factor(s) might regulate *prdx-2* expression in these neurons. Consistently, several transcription factors were reported to bind to the *prdx-2* promoter by ChIP-seq [42] (S3 Fig).

We conclude that *skn-1* accomplishes a cell autonomous regulation of *prdx-2* in the intestine, but that this regulation may not occur in neurons. Taken together, our data suggest that PRDX-2 might exert a different function depending on the cell type: H_2_O_2_ detoxification in the foregut, triggered by SKN-1, *vs.* H_2_O_2_ perception or signaling in I2 neurons.

### PHA neurons respond to H_2_O_2_ in a prdx-2-dependent manner

As PRDX-2 is expressed in other neurons than I2s, we asked whether PRDX-2 endows these neurons with oxidative stress sensing properties. We focused on PHA tail neurons, as they belong to the phasmid sensory sensilla and respond to many noxious stimuli [12]. We monitored PHA neuron activation by imaging calcium using the GCaMP3 fluorescent sensor [44] expressed under the *flp-15* promoter, specific to I2 and PHA neurons [45].

To image calcium fluxes in neurons, L4 animals were trapped in microfluidic chambers and exposed to H_2_O_2_ (Fig 3A, S4 Fig), and their response was followed in 4D spinning-disc ultrafast acquisitions. This experimental setup, combined with semi-automated image analyses to quantify the mean fluorescence in neurons over time (Supplementary Methods), confirmed the activation of I2 neurons upon a 1mM H_2_O_2_ treatment (Fig 3B,D, Movie 1, S5 Fig), similar to that observed upon exposure to H_2_O_2_ vapor (*ie.* 8.82M,[8]). As PRDX-2 reproducibly shows an asymmetric expression level in I2L and I2R (Fig 1B,D and detailed description in [73]), each neuron was scored individually to take into account a putative left-right effect. However, this did not impact neuron response as no significant difference in the normalized response of left and right neurons was noticed (Fig 3B,L). Importantly, we observed that PHA neurons respond to 1mM H_2_O_2_ comparably to I2 neurons (Fig 3C,D,H, Movie 2, S5 Fig). We then investigated whether PRDX-2 function was necessary for PHA response, using a strong *prdx-2* loss of function mutant. In agreement with previous work [8], I2 neurons in *prdx-2(gk169)* mutants failed to respond to 1mM H_2_O_2_ (Fig 3E,F,H, Movie 3, S5-S6 Figs). Interestingly, we uncovered that PHA neuron response was severely impaired in *prdx-2* mutants, although not completely abolished as in I2s (Fig 3E,G,H, Movie 4, S5-S6 Figs). From these observations, we conclude that both I2s and PHAs responses to oxidative stress require the function of the peroxiredoxin PRDX-2, but the residual response of PHA neurons suggests a lesser requirement of the antioxidant in tail neurons.

**Figure 3.**
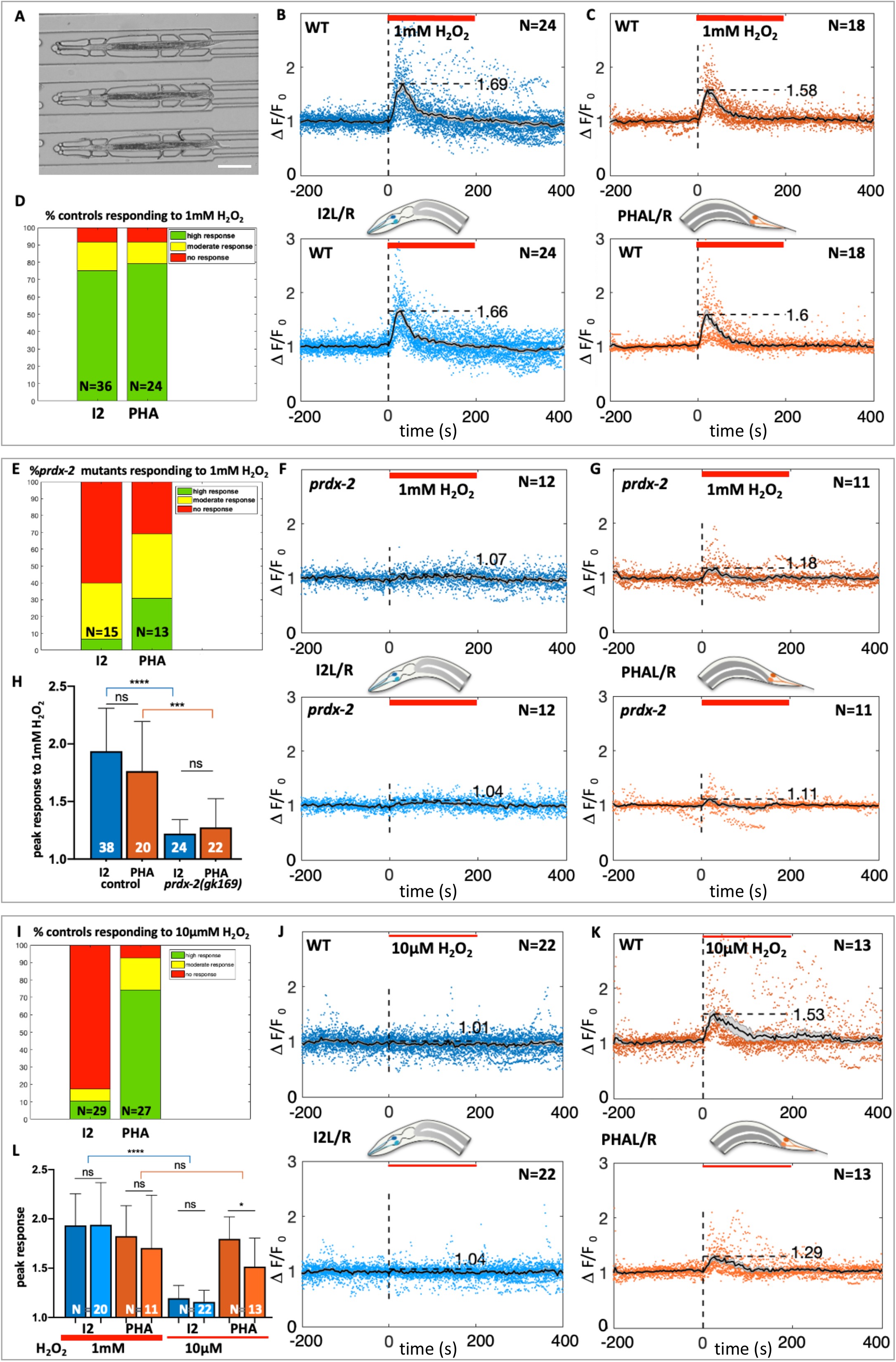
I2 and PHA neurons both respond to 1mM H_2_O_2_ in a *prdx-2*-dependent manner, but only PHA neurons respond to 10μM H_2_O_2_. (A) Low-magnification DIC picture of L4 animals trapped in the microfluidic device used in all neuron recordings experiments. (B,C,F,G,J,K) Calcium response (visualized by GCaMP3) in I2R/I2L (B,F,J) and in PHAL/PHAR neurons (C,G,K) following H_2_O_2_ treatment (indicated by the red bar). Average curves show normalized neuron responses (time in seconds), in both left and right neurons (top and bottom curves). N, number of movies quantified for each genotype, in wild-type (B,C,J,K) and in *prdx-2* mutants (F,G). (D,E,I) Bar graph showing the fraction of animals responding to the H_2_O_2_ stimulation in all experiments, classified as high (green), moderate (yellow) or absent (red) responses (see Methods). N, number of movies analyzed. (H,L) Quantification of the calcium response to H_2_O_2_ in I2 and PHA neurons in controls and in *prdx-2* mutants at 1mM (H), and in I2L/R and PHAL/R controls, at 1mM and 10μM H_2_O_2_ (L). Bars represent the mean (N, number of movies) and error bars SD. ns, not significant, p>0.05; *p<0.05; ***p<0.001; ****p<0.0001 (Mann Whitney and Kruskal-Wallis tests). See corresponding Movies 1-6 and S5 Fig.

### I2s and PHAs show differences in H_2_O_2_ sensitivity and in receptors involved

Given the putative role of PRDX-2 as an H_2_O_2_ sensor, the fact that there was a slight difference in PRDX-2 activity requirement in I2s and PHAs prompted us to analyze whether the head and tail neurons share the same sensitivity to H_2_O_2_. We thus tested whether I2 and PHA neurons exhibit a response to the very mild dose of 10μM H_2_O_2_, a dose which induces a less penetrant pharyngeal pumping inhibition (≈ 35% of animals) than that observed at 1mM (≈ 90% of animals, [8]). In our experiments, whereas I2 neurons failed to be activated in most animals at 10μM H_2_O_2_ (25/29, Fig 3I,J,L, Movie 5, S5 Fig), PHA neurons responded in the vast majority of animals (25/27, Fig 3I), in a similar manner than at 1mM H_2_O_2_ (Fig 3K,L, Movie 6, S5 Fig). In addition, PHA neurons response to 10μM H_2_O_2_ depends on PRDX-2, as *prdx-2* mutants PHA neurons all fail to respond to this low dose (13/13, Movie 7, S5-S6 Figs). The requirement of PRDX-2 for PHA response to micromolar H_2_O_2_ suggests that PRDX-2 is unlikely to transmit the H_2_O_2_ signal under its hyperoxidized form. Taken together, we conclude that PHA neurons are more sensitive to low doses of H_2_O_2_ than I2 neurons.

The difference in sensitivity between I2 and PHA neurons may come from distinct molecular mechanisms. To explore this possibility, we first tested which receptors are required in I2s and in PHAs for H_2_O_2_ perception. We focused on photoreceptors as photosensation is likely to involve the generation of ROS [8,46]. In *C. elegans*, light sensing relies on unusual gustatory G-protein-coupled receptors (GPCRs) related to vertebrate photoreceptors: the two nematode closest paralogs LITE-1 and GUR-3 mediate photosensation in ASJ and ASH neurons [46–48], and light and H_2_O_2_ sensing in I2 neurons [8]. We investigated whether these two receptors are differentially localized in I2 and PHA neurons. We generated a knock-in GUR-3::GFP line, which revealed that GUR-3 is solely expressed in I2 and I4 photosensory neurons (Fig 4A), as previously reported using episomal expression [8]. Such a restricted pattern deeply contrasts with the broad expression domain of LITE-1, which includes phasmid neurons (PHA and PHB), but not I2 neurons, as shown using both translational and transcriptional reporters [8].

**Figure 4.**
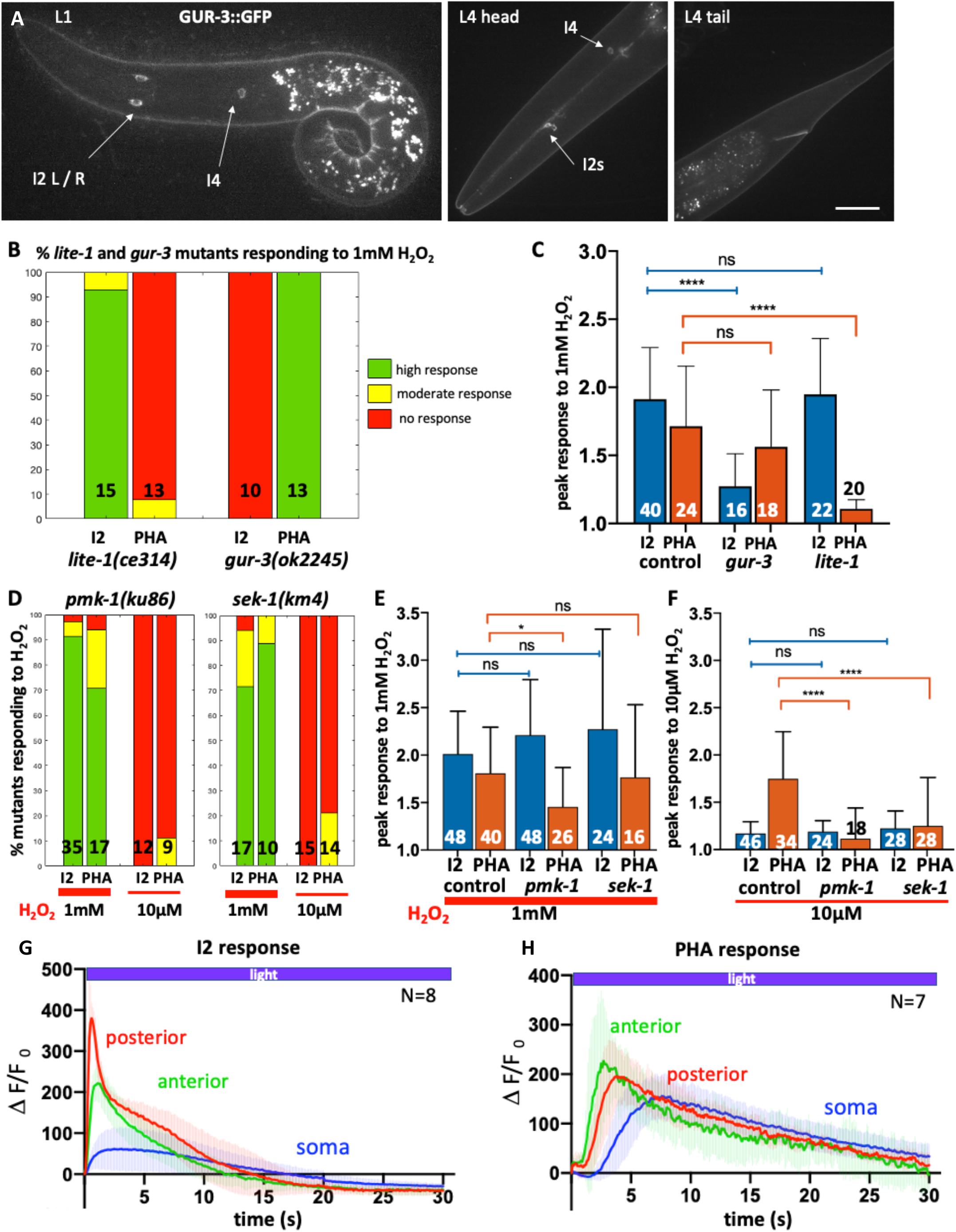
H_2_O_2_ response involves different receptors and transducers in I2 and PHA neurons. (A) Spinning-disc confocal projections of a representative GUR-3::GFP knock-in animal, at L1 (left) and L4 stages (right). The GUR-3::GFP signal is detected in I2 neurons and in a single I4 neurons (head panel), but not in PHA/PHB neurons (tail panel). Bar, 10μm. (B,D) Bar graph of the fraction of *lite-1, gur-3, pmk-1* and *sek-1* mutants responding to the H_2_O_2_ stimulation, classified as high (green), moderate (yellow) or absent (red) responses. N, number of movies analyzed. (C,E,F) Quantification of the calcium response to 1mM H_2_O_2_ in I2 and PHA neurons in controls and in *gur-3* and *lite-1* mutants (C), and in *pmk-1* and *sek-1* mutants at 1mM (E) and at 10μM H_2_O_2_ (F). Bars represent the mean (number of movies indicated) and error bars SD; ns, not significant, p>0.05; *p<0.05; ****p<0.0001 (ANOVA and Kruskal-Wallis tests). See corresponding Movies 8-19 and S7-S10 Figs. (G,H) Blue light triggered different calcium fluxes in I2 and PHA neurons in the three regions analyzed; anterior neurite (green), posterior neurite (red) and soma (blue). The curves represent the mean and shaded error bars indicate SD. See Movies 20-22.

These differential localizations prompted us to inquire whether mutants in these receptors were still able to trigger a response to 1mM H_2_O_2_ in I2 and PHA neurons. In *gur-3(ok2245)* mutants, only PHA neurons were able to respond to 1mM H_2_O_2_ (Fig 4B,C, Movies 8,9, S7-S8 Figs), providing evidence that GUR-3 function is not essential in PHA neurons for H_2_O_2_ sensing. In contrast, *lite-1(ce314)* mutants showed a reciprocal response, with only I2 neurons responding to 1mM H_2_O_2_ (Fig 4B,C, Movies 10,11, S7-S8 Figs). In conclusion, H_2_O_2_ likely activates I2 neurons via *gur-3* and PHA neurons via its paralog *lite-1*, respectively. Interestingly, these observations might explain the previously reported impaired avoidance response of *lite-1* mutants to 1mM H_2_O_2_ [8], as PHA neurons are involved in escape behavior [13].

### PMK-1 and SEK-1 are required for PHA neurons response to micromolar H_2_O_2_ but are dispensable in I2 neurons

Since H_2_O_2_ perception in I2 and PHA neurons involves different receptors and requires the function of PRDX-2 in both cases, we wondered what type of signaling occurs downstream PRDX-2 to trigger neuronal activation upon H_2_O_2_ stimulation.

As mentioned above, studies in various models have reported that peroxiredoxins can modulate the p38/MAPK signaling pathway to influence cellular decisions, notably in drosophila and mammalian cells [34]. In *C. elegans*, specifically, the activation of this PRDX-2-PMK-1/p38MAPK cascade allows micromolar doses of H_2_O_2_ to potentiate the ASH neuron sensory behavior to glycerol [3]. Therefore, we investigated whether the p38/MAPK pathway could be involved in H_2_O_2_ sensing by analyzing the responses of *pmk-1* (p38/MAPK) and *sek-1* (MAPKK) mutants in I2 and PHA neurons. As the strongest allele *pmk-1(ok811)* is homozygous lethal, we had to use the hypomorphic *pmk-1(km25)* viable mutant, which carries a N-terminal deletion [49]. In *pmk-1(km25)* mutants, while I2 neurons responded normally to 1mM H_2_O_2_, PHA neurons showed a slightly milder response to 1mM H_2_O_2_ compared to controls (Fig 4D,E, Movies 12,13, S9 Fig). In *sek-1(km4)* MAPKK mutants, we observed comparable responses of I2 and PHA neurons to 1mM H_2_O_2_ to that of controls (Fig 4D,E, Movies 16,17, S10 Fig). We conclude that for 1mM H_2_O_2_ sensing, the p38/MAPK pathway is dispensable at least in I2 neurons, but may play some role in PHA neurons.

In ASH neurons, the PRDX-2-mediated activation of the p38/PMK-1 cascade induces a potentiation of their sensory behavior [3]. Therefore, we asked whether the p38/PMK-1 pathway could similarly promote PHA higher sensitivity to H_2_O_2_. We examined PHA neurons response to 10μM H_2_O_2_ in *pmk-1* (MAPK) and in *sek-1* (MAPKK) mutants. Strikingly, both mutants displayed a very similar phenotype, with an abolished response of PHA neurons to micromolar doses of H_2_O_2_ observed in *pmk-1* (Fig 4D,F, Movie 15, S9 Fig) and in *sek-1* mutants (Fig 4D,F, Movie 19, S10 Fig). In I2 neurons, as in controls, both *pmk-1* and *sek-1* failed to respond to 10μM H_2_O_2_ (Fig 4 D,F, Movies 14, 18, S9-S10 Figs), as expected. Taken together, this suggests that the p38/MAPK pathway would be specifically required for PHA neurons hypersensitivity to H_2_O_2_, but dispensable in I2 neurons.

### PHA neurons are photosensory neurons like ASH neurons

Light sensing has been reported for ASJ, ASH and I2 neurons and require either the LITE-1 or the GUR-3 receptor, respectively [8,46–48]. As PHAs neurons require LITE-1 to respond to H_2_O_2_ (Fig 4B,C, Movie 11, S7 Fig), we asked whether they could respond to light. To test this, we monitored calcium transients in three neuronal compartments using the GCaMP strain upon stimulating neurons with blue light, as previously done for I2 or ASH neurons [8,46]. Interestingly, we found that all regions of PHA neurons responded to light, displaying a different response profile than those of I2 neurons (Movie 22): PHA soma showed a stronger and longer response than I2 soma; PHA posterior neurite responded much slower that in I2 where it exhibits the fastest and strongest response peak, and the anterior neurite also had a slower recovery than in I2 (Fig 4G,H, Movies 20,21). Overall, while I2 neurons exhibit a fast photoresponse within 10-15s, PHA neurons photoresponse requires twice longer to return to steady state (approx. 30s). Strikingly, we noticed that PHA neurons profile is highly reminiscent of that reported in ASH neurons photoresponse [46].

It has been proposed that light sensing may involve intracellular H_2_O_2_ release and peroxiredoxin signaling in both nematodes and yeast [7,8]. To test whether PHA response to light requires the peroxiredoxin PRDX-2, we analyzed light sensing in *prdx-2* mutants. As in H_2_O_2_ sensing (Fig 3 E,F,H, Movie 3), we observed that I2 response to light strictly depends on *prdx-2* (15/15, Movie 23, S11 Fig). Surprisingly, we found that PHA photoresponse does not require PRDX-2, as *prdx-2* mutants were still able to respond to light (17/20, Movie 24, S11 Fig). We conclude that light sensing does not involve the same mechanisms in I2 and in PHA neurons, downstream the photoreceptors.

## Discussion

### H_2_O_2_ sensing in head and tail neurons relies on different mechanisms

Here, we describe how two pairs of sensory neurons located in the head and the tail of *C. elegans*, namely I2s and PHAs, contribute to exogenous H_2_O_2_ and light sensing. Compared to previous reports, our study relies on a PRDX-2::GFP knock-in line more closely reflecting endogenous expression level, in comparison to overexpression often observed with extrachromosomal arrays. While classical methods do not enable precise control of the environment such as application of a stress, we carried out neuron response experiments using the microfluidic technology, allowing live imaging of immobilized animals upon simultaneous exposure to a controlled oxidative stress.

We found that PHA tail neurons can elicit a response to a micromolar range of H_2_O_2_, whereas I2 head neurons cannot, suggesting that distinct molecular mechanisms may account for this difference. Accordingly, while the peroxiredoxin PRDX-2 is essential for H_2_O_2_ sensing in both I2 and PHA neurons, a different transmembrane receptor is required to transduce the signal: I2 neurons use the gustatory receptor GUR-3, while PHA neurons would require its paralogue LITE-1. Another difference lies in the p38/MAPK activity requirement: while dispensable in I2 neurons, it would be specifically required in PHA neurons to confer their hypersensitivity to H_2_O_2_ (lack of response of PHA neurons to 10μM H_2_O_2_ in *sek-1*/MAPKK and in *pmk-1/MAPK* mutants, Fig 4). Finally, although nematodes were known for a long time to avoid light when the light pulse was applied on their tail [47], our work provide the first evidence, to our knowledge, that PHA tail neurons act as photoreceptor cells.

Overall, our data are consistent with previous findings unveiling the existence of two distinct modes of response to oxidative stress in *C. elegans*: a direct response in peripheral tissues such as the gut, and a neuronally-regulated response relying on synaptic transmission [50]. Specifically, we uncovered that a harsh oxidative stress (10mM H_2_O_2_) triggers PRDX-2 induction in the anterior gut and in the EPC (Figs 1-2), while lower doses of H_2_O_2_ triggers either I2 and PHA neurons activation (1mM), or only PHA neurons activation (10μM) (Fig 3). Interestingly, both types of response involve the peroxiredoxin PRDX-2, which behaves differently in the two cellular contexts: PRDX-2 would be cell-autonomously induced by SKN-1 in the intestine, but likely not in neurons (Fig 2). Therefore, we propose that PRDX-2 might act as a peroxidase in the gut, as proposed [27], whereas it could function as a H_2_O_2_-signaling molecule in neurons, as suggested for I2 neurons [8]. Importantly, H_2_O_2_ response in I2 and PHA neurons requires the joint function of PRDX-2 and a receptor, as each mutant individually cannot respond (Figs 3-4). In addition, former rescue experiments indicated that light response requires the activity of PRDX-2 and GUR-3 specifically in I2 neurons [8]. Based on all these data and recent studies shedding light on how H_2_O_2_ is sensed in plant and animal cells [34,51], we propose new hypotheses regarding H_2_O_2_ signaling in neurons which are depicted in Fig 5 and described below, to integrate our findings with recent results from the literature.

**Figure 5.**
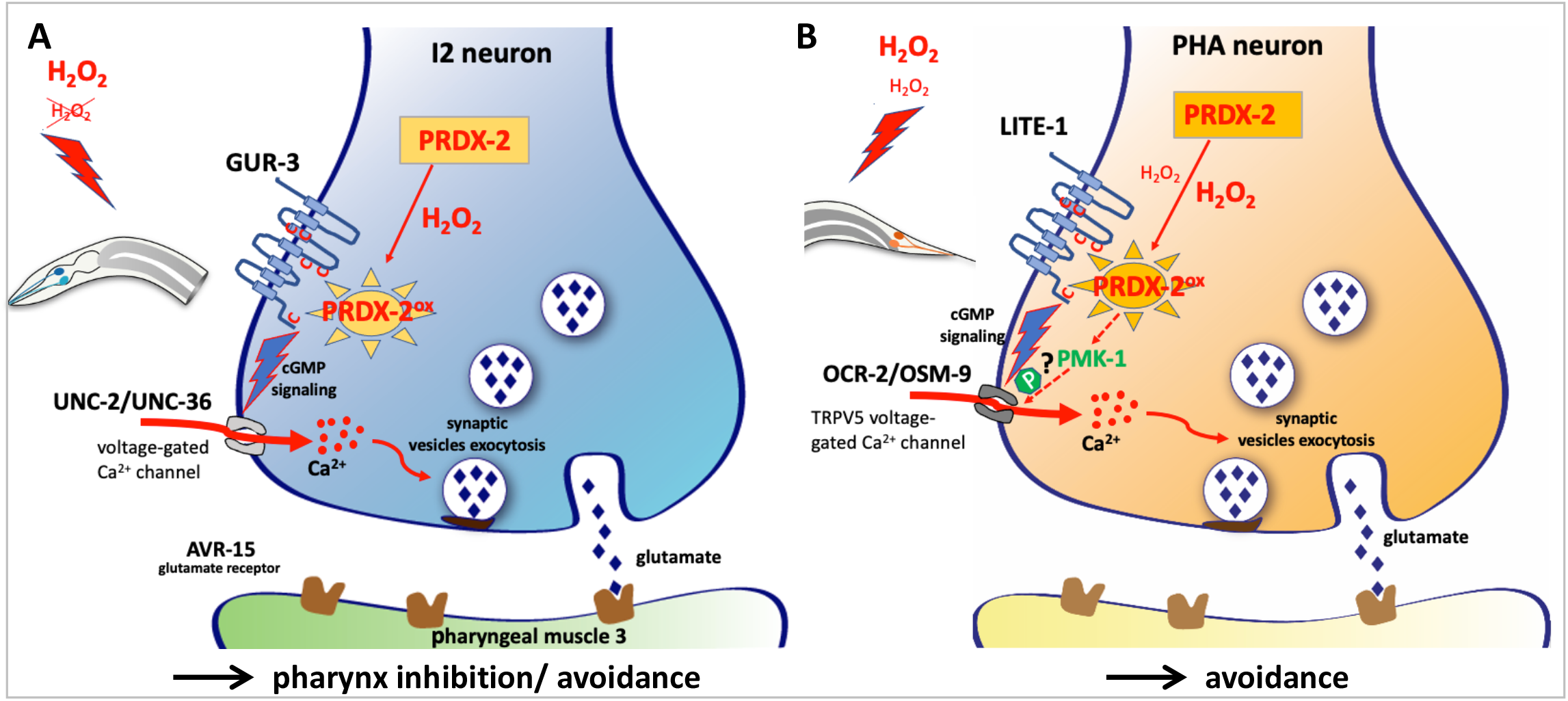
Hypothetical model of H_2_O_2_ sensing and signaling in *C. elegans* I2 and PHA neurons based on this and on previous work. (A-B) Hypothetical model of H_2_O_2_-induced neuronal activation, based on our data and on previous studies. Schematic drawing of a presynaptic button in I2 (A) and PHA (B) neurons, illustrating the presumptive H_2_O_2_-PRDX-2-mediated neuronal activation in both cases (modelled in S12 Fig). High doses of H_2_O_2_ (1mM) are sensed by both neurons, but only PHA neurons respond to 10μM H_2_O_2_, as illustrated in red. In this model, H_2_O_2_ would freely diffuse through the neuron plasma membrane and oxidize PRDX-2, presumably leading to LITE-1 or GUR-3 activation. Receptor activation is likely relayed by cGMP signaling, resulting in the opening of voltage-gated calcium channels (in grey) and neurotransmitter release (glutamate in both cases), triggering an adapted response. In PHA neurons, the PMK-1/p38MAPK pathway is additionally required to promote neuronal response to micromolar doses of H_2_O_2_, potentially through OSM-9 phosphorylation, as observed in ASH neurons [3]. See discussion for further details and bibliographic references.

### A presumptive model of H_2_O_2_ sensing in C. elegans neurons

In both I2 and PHA neurons, we favor the hypothesis that cytosolic PRDX-2 rather than the transmembrane receptor would be the neuronal H_2_O_2_ sensor, based on the following observations: i) in many cases, H_2_O_2_ signaling is mediated by oxidation of cysteines in redox-regulated proteins [52]. Alternatively, redox signaling often relies on peroxiredoxins acting as sensor and transducer of H_2_O_2_ signal [53], as thiol modifications would be much faster when catalyzed by peroxiredoxins [54–56], due to their abundance and their high reactivity to H_2_O_2_ [57]. Here, the striking abundance of PRDX-2 in PHA, and especially in I2 neurons reinforces this idea, as well as the defective response of *prdx-2* mutants to both doses of H_2_O_2_ tested (Fig 3, S6 Fig). Consequently, we propose that H_2_O_2_ signaling in I2 and PHA neurons would involve redox signaling through PRDX-2. ii) Concerning receptor topology, GUR-3 and LITE-1 present a higher conservation in their intracellular domains, with conserved cysteines only found intracellularly and in a transmembrane domain (S12A Fig). This structure is more reminiscent of that of the vertebrate transmembrane protein GDE2, which is activated intracellularly by the Prdx1 peroxiredoxin [58], than of the plant H_2_O_2_ sensor, the HCPA1 receptor, whose activation involves direct oxidation of extracellular cysteines [51]. Here, GUR-3 and LITE-1 receptor topology does not support the hypothesis of a direct oxidation by H_2_O_2_ on the extracellular domain, but rather suggests intracellular signaling as in the case of GDE2 (S12B,C Fig). We thus propose a scenario in which H_2_O_2_ would diffuse through the neuron plasma membrane and oxidize PRDX-2. Oxidized PRDX-2 or its disulfide form (PRDX-2^ox/S-S^) would in turn react with the cysteines of GUR-3 or LITE-1, triggering receptor activation, and I2 or PHA neuron response (Fig 5, S12 Fig). These hypotheses will require future experimental validation.

Neuronal response involves the opening of calcium channels, which are different in I2 and PHA neurons: I2 activation depends on UNC-2 and UNC-36 voltage-gated calcium channels [9], while PHA neurons require the cyclic nucleotide-gated channel TAX-4 and the vertebrate TRPV5 nematode equivalent OSM-9 [12]. Of note, the other TRPV5 channel subunit OCR-2 may also act in PHAs, as it is expressed in PHAs (S13 Fig, [59]) and it functions together with OSM-9 [60]. In conclusion, although closely related and both able to sense H_2_O_2_ and light, GUR-3 and LITE-1 receptor signaling likely involves different downstream transducers. Except for PRDX-2 requirement, I2 and PHA neurons would use distinct molecular pathways to transduce H_2_O_2_ response.

### A striking parallel between PHA and ASH neurons

Our data uncover a high sensitivity of PHA neurons to micromolar doses of H_2_O_2_ that is not seen in I2 neurons and requires PRDX-2 and p38/MAPK activity. Strikingly, micromolar doses of H_2_O_2_ were also reported to activate the p38/MAPK pathway via PRDX-2 in ASH neurons, leading to the phosphorylation of the OSM-9 TRPV sensory channel, thereby increasing its sensitivity [3]. Although our study and the latter do not elucidate how PRDX-2 triggers PMK-1 activation, recent evidence sheds light on this process: in both mammalian and drosophila cells, H_2_O_2_ induces transient disulfide-linked conjugates between the MAP3K and a typical 2-Cys peroxiredoxin [34]. Similarly, in *C. elegans*, PRDX-2 could activate the MAPKKK NSY-1, as NSY-1 function is required in ASH neuron [3], and may be expressed in PHAs *(unidentified cells of the tail, D. Moerman, WormBase).* Based on these observations and on the fact that OSM-9 is expressed [61] and likely required in PHA neurons [12], it is possible that PMK-1/p38MAPK activation might increase PHA neurons’ sensitivity to H_2_O_2_ through the downstream phosphorylation of OSM-9, via AKT-1, as in ASH neurons (Fig 5). Alternatively, by analogy with recent work showing that DLK/p38MAPK signaling controls LITE-1 stability in ASH neurons, PMK-1 could also control LITE-1 turnover by RAB-5-mediated endocytosis [62]. These assumptions, which will necessitate further experimental testing, are supported by the observation that ASHL/R and PHAL/R are both descendants of ABplp or ABprp in the nematode cell lineage, and hence may express similar sets of genes following lineage-specific priming [63].

The fact that PHA, PHB and ASH were found in the same neuron cluster [64] confirms that they share similar molecular signatures. To illustrate the parallel between ASH and PHA polymodal nociceptors, we retrieved the set of genes expressed in these neurons [64], and analysed which ones were specifically enriched in these neurons in comparison to all other neurons (S13 Fig, data in S1-S3 Tables). This analysis revealed that ASH and PHA not only express many genes specific to ciliated neurons, as expected (*eg*. *che-3, nphp-4, che-11, ifta-1*, C33A12.4 and R102.2, [65]), but also a number of common receptors (*eg. ocr-2, osm-9, ida-1, casy-1, ptp-3* or *pdfr-1)*, although some of them are more enriched in ASH *(dop-2, nrx-1, snt-5, sue-1).* Intriguingly, their neuropeptide profile appears different for the two neuron pairs (PHA enriched in *flp-7, flp-4, flp-16, nlp-1 ins-18*, while ASH strongly express *flp-13* and *npr* genes), suggesting that beyond their functional similarity, ASH and PHA may trigger different types of intercellular communication. Finally, this analysis highlights the fact that several genes with unknown function are highly enriched in both ASH and PHA (*eg*. F27C1.11, W05F2.7, *tos-1, cab-1)*, pointing to their potential role in neuronal function.

Taken together, our analyses highlight the common features shared between PHA and ASH polymodal nociceptive neurons, as formerly noticed [13], since they both: i) display a higher sensitivity dependent on the p38/MAPK pathway (Fig 4D-F and [3]), ii) exhibit a similar response profile to light (Fig 4H and [46]), iii) require the photoreceptor LITE-1 for lightsensing [46] or H_2_O_2_ sensing (Fig 4BC), iv) trigger avoidance [13], and v) share a close molecular signature (S13 Fig and [64]). A recent report indicates that ASJ neurons are also required for H_2_O_2_ avoidance [6], and unravels the behavioral mechanisms allowing *C. elegans* to find a suitable niche, owing to the interplay between H_2_O_2_ and bacteria in its environment. Overall, these observations led us to propose that nematodes might integrate the environmental redox signals from at least four different pairs of neurons (I2, ASH, PHA and ASJ) in order to trigger an appropriate dose-dependent physiological response.

### GUR-3 and LITE-1 receptors mediate both light and H_2_O_2_ sensing

In yeast, light sensing relies on the peroxisomal oxidase Pox1 which triggers light-dependent H_2_O_2_ formation, the latter being sensed by the Tsa1 peroxiredoxin and transduced to thioredoxin for subsequent signaling [7]. Nematodes, unlike yeast, require a photoreceptor in addition to the antioxidant for lightsensation: GUR-3 in I2 neurons [8], and LITE-1 in ASJ and ASH neurons [46,48]. Here we showed that both I2 and PHA respond to light, but with a different profile. Despite its unusual membrane topology, LITE-1 has been shown to encode a bona fide photoreceptor, whose photoabsorption depends on its conformation [66]. However, whether light directly activate the neuron photoreceptor or triggers intracellular H_2_O_2_ release and signaling is still unclear. In agreement with the fact that H_2_O_2_ inhibits LITE-1 photoabsorption *in vitro* [66], it has been shown that a H_2_O_2_ pretreatment reduces LITE-1-mediated photoresponse in ASH neurons [46]. Consistent with these reports, our data illustrate that GUR-3 and LITE-1 have a dual function in both light and H_2_O_2_ sensing. In I2 neurons, as in yeast, redox signaling could be involved in transducing the light signal, as PRDX-2 is strictly required for light sensing (S11 Fig, [8]). In contrast, PHA neurons can respond to light without PRDX-2, indicating that the LITE-1 photoreceptor tranduces the light signal without a redox relay. This difference could explain our observation that I2 neurons respond faster to light than PHA and ASH neurons (Fig 4 G,H and [46]). Whether this difference strictly depends on the photoreceptor (LITE-1 in PHA, ASH *vs* GUR-3 in I2) and/or on its downstream signaling cascade is an open question.

Finally, it is noteworthy that I2 and PHA neurons rely on peroxiredoxin for H_2_O_2_ and/or light sensing, while ASJ and ASH rely on thioredoxin: TRX-1 is required for LITE-1- dependent photosensation in ASH neurons [46], and is expressed in ASJ photosensory neurons [67], which respond to H_2_O_2_ [6]. Altogether, this further underlines the importance of a redox signaling relay involving antioxidants of the peroxiredoxic cycle in nematode photosensory neurons.

In conclusion, our work illustrates that nematodes can sense various concentrations of H_2_O_2_ through sensory neurons located in the head and the tail, using either partially different (I2 and PHA), or similar molecular mechanisms (ASH and PHA). This set of neurons also confer light sensing to the nematode with a distinct speed in the response (fast in I2, slow in PHA and ASH, [46]). While I2 neurons seem more specialized in sensing oxidative stress [8,10], PHA and ASH can detect many other stimuli [12], but all of them can trigger avoidance. To tackle the complex question of how these neuronal inputs translates into behavior, a recent method was developed [68], allowing the simultaneous recording of behavioral and neural responses of *C. elegans* to salt concentrations changes. These observations may help uncover whether and how inputs from head and tail oxidative stress sensory neurons integrate to allow nematodes to quickly and appropriately react to a change in the environment.

## Experimental procedures

### Generation of plasmids and transgenic strains by CRISPR/Cas9-mediated genome editing

*C. elegans* strains (listed in Supplementary Materials) were maintained as described [69]. PRDX-2::GFP knock-in strain was generated by CRISPR/Cas9-mediated genome editing, using a DNA plasmid-based repair template strategy [37]. For both PRDX-2 and GUR-3 knockins, a C-terminal GFP fusion was generated, comprising a flexible linker between the coding region and GFP to allow correct folding of the fusion protein. A combined small guide-RNA/repair template plasmid was built using the SAP Trap strategy [70]. Phusion DNA polymerase was used to amplify by PCR 5’ and 3’ *prdx-2* and *gur-3* homology arms (HAs) from N2 genomic DNA, using primers containing SapI restriction sites and silent mutations to prevent Cas9 re-cleavage (all primers sequences listed in Supplementary Materials). After purification, 5’ and 3’ HAs and sgRNA oligonucleotides were assembled into the destination vector (pMLS256), together with the flexible linker (from pMLS287), GFP and the *unc-119* rescuing element (from pMLS252), in a single SapI restriction-ligation reaction, as described [70]. Prior to transformation in DH5α cells, a sabotage restriction was performed with SpeI to digest empty destination vectors but not the desired assembly constructs, which were subsequentially verified by restriction digest analysis and sequencing. All plasmids used for injection were purified using a DNA Miniprep Kit (PureLink, Invitrogen), or a DNA midiprep kit (Macherey Nagel). For PRDX-2::GFP knock-in, a plasmid mix containing combined sgRNA/repair template plasmid (50 ng/μl), Cas9-encoding pSJ858 (25ng/μl) and co-injection markers (pCFJ90 at 2.5ng/μl; pCFJ104 at 5ng/μl, and pGH8 at 5ng/μl) was injected in the germline of *unc-119(ed3)* animals [37]. For GUR-3::GFP knock-in, an injection mix containing purified Cas9 protein (IDT) associated with tracrRNA and crRNA (guide RNA), GUR-3 repair template, and co-injection markers was injected in *unc-119(ed3)* animals, according to IDT online protocols for *C. elegans*. Plates containing 2/3 injected F0 animals were starved, chunked on fresh plates, for candidates screening (attested by the presence of wild-type non fluorescent animals). Knock-in events were validated by PCR on homozygous lysed worms (QuantaBio AccuStart II GelTrack PCR SuperMix), using primers annealing in the inserted sequence and in an adjacent genomic region not included in the repair template. The PRDX-2::GFP strain was outcrossed 5 times to N2 wild-types.

### RNA interference

RNAi experiments were performed by feeding using the Ahringer-MRC feeding library [71]. Animals fed with the empty vector L4440 served as a negative control. The efficiency of each RNAi experiment was assessed by adding an internal positive control, *zyg-9(RNAi)*, which induces embryonic lethality.

### Spinning-disk confocal microscopy acquisitions and fluorescence intensity measurements

For live imaging, animals were anesthetized in M9 containing 1mM levamisole and mounted between slide and coverslip on 3% agarose pads. Synchronized L4 animals were treated for 30min in a 96-well flat bottom plate, in 50μl of M9 containing 1mM or 10mM H_2_O_2_. Treated animals were transferred using a siliconized tip on a freshly seeded plate to recover, and imaged 1h30 to 2h later. Spinning-disk confocal imaging was performed on a system composed of an inverted DMI8 Leica microscope, a Yokogawa CSUW1 head, an Orca Flash 4.0 camera (2048*2018 pixels) piloted by the Metamorph software. Objective used were oil-immersion 40X (HC PL APO, NA 1.3) or 63X (HCX PL APO Lambda blue, NA 1.4). The temperature of the microscopy room was maintained at 20°C for all experiments. Z-stacks of various body regions were acquired with a constant exposure time and a constant laser power in all experiments. Maximum intensity projections were used to generate the images shown. Fluorescence intensity measurements in int1, I2 and EPC cells were performed using the Fiji software, by manually drawing a region of interest (ROI) around the cell (int1, EPC), or applying a threshold (I2 neurons), background was subtracted and average pixel intensity was quantified.

### Microfabrication and microfluidic chip preparation

The microfluidic chip original design was inspired by the the Wormspa [72], but pillars distances were adapted to trap L4 animals, and multiple series of traps were included to increase the number of experiments per chip (S4 Fig). A master mold was made by standard soft photolithography processes by spin-coating a 25μm layer of SU-8 2025 (Microchem, USA) photoresist at 2700 rpm for 30sec on a 3” wafer (Neyco, FRANCE). Then, we used a soft bake f 7min at 95°C on hot plates (VWR) followed by a UV 365nm exposure at 160 mJ/cm^2^ with a mask aligner (UV-KUB3 Kloé^®^, FRANCE). Finally, a post-exposure baking identical to the soft bake was performed before development with SU-8 developer (Microchem, USA). Then, the wafer was baked at 150°C for 15min to anneal potential cracks and strengthen the adhesion of the resist to the wafer. Finally, the master mold was treated with chlorotrimethylsilane to passivate the surface.

Worm microchannels were cast by curing PDMS (Sylgard 184,10:1 mixing ratio), covalently bound to a 24 × 50 mm coverslip after plasma surface activation (Diener, Germany), and incubated 20min at 60°C for optimal adhesion. The chip was perfused with filtrated M9 solution through the medium inlet using a peristaltic pump (Ismatec), until complete removal of air bubbles. Worm loading was performed with the pump set at a low flow rate (<30 μl/min), through a distinct inlet (S4 Fig): 10-15 young L4 animals (synchronized by bleaching 48h prior to each experiment) were picked in a siliconized Eppendorf tube (Sigmacote SL2, Sigma Aldrich) containing M9, and perfused into the traps. The loaded chip was carried to the microscope while being still connected to the pump by a gravity flow (preventing animals to escape) until the microfluidic chip was installed on the microscope stage.

### Calcium imaging

I2 and PHA neuronal response was monitored using the calcium sensor GCaMP3 expressed under the *flp-15* promoter as in [8]. To image H_2_O_2_ response, young L4 animals trapped in microfluidic chips were imaged using the confocal spinning disc system described above with the 20X air objective (HC PL APO CS2, NA 0.75). The microfluidic chip allowed the simultaneous recording of up to 3 animals per experiment (S4 Fig). Z-stacks of 10-15 images (10 μm spacing) were acquired every 2s (using the stream Z mode), for 350 time points. Exposure time was 50ms and laser power set on 40%. The device was perfused with M9 medium throughout the experiment using a peristaltic pump set at 80 μl/min, and H_2_O_2_ was perfused (at 10μM or 1mM in M9) for 100 time points (3min20s) after an initial recording of 35-45 time points. Movies were computationally projected using MetaMorph, and data processing (including movie registration, neuron segmentation and tracking over time) was conducted with a custom-developed Matlab program detailed below and available at https://github.com/gcharvin/viewworm (a user guide is provided in Supplementary Methods).

To image light response in I2 and PHA neurons, L4 worms were mounted on a slide covered with 3% agarose pads in M9 supplemented with 1mM levamisole. Video-recordings were performed on the spinning-disc microscope using the 40X oil objective. Animals were exposed to blue light (485nm) while their neuronal response was simultaneously recorded in stream mode (10 frames/sec, single Z, 100ms exposure, laser 100%) for 30sec.

### Calcium response analyses

For H_2_O_2_ response analyses, sequences of images were spatially realigned with respect to the first image of the timeseries in order to limit the apparent motion of the worm in the trap and ease the tracking of neurons of interest. This image registration process was performed using standard 2D image cross-correlation by taking the first image as a reference. Then, we used a machine-learning algorithm (based on a decision tree) to segment pixels in the fluorescent images. For this, we took a series of image transforms (gaussian, median, range filters) as descriptors for the classifier, and we trained the model on typically 10 frames before applying the result to the rest of the time series. This segmentation method appeared to be superior to simple image thresholding, which is inadequate when dealing with fluorescent signals that vary both in time and space (the brightness of two neurons is quite different). Next to the segmentation procedure, we tracked the identified neurons using distance minimization, and we quantified the mean fluorescence signal in each neuron over time. Last, fluorescence data corresponding to individual animals were pooled after synchronization from the time of exposure to H_2_O_2_ and signal normalization. This image analysis pipeline is available at https://github.com/gcharvin/viewworm and a tutorial for use is included in Supplementary Methods. As some movies could not be quantified due to uncontrolled animal movements, a visual classification of neuronal responses (high, moderate, absent) was made by comparison to successfully tracked movies.

For light response analyses in I2 and PHA neurons, the same image processing pipeline as in [8] was used, except that ROI were manually drawn in Fiji.

### Statistical analyses

For pairwise comparisons of data sets with a normal distribution (or N>30), p-values were calculated using an unpaired two-tailed Student test, and the Welch correction was applied when samples variance was not homogeneous. When distributions were not normal (Anderson-Darling and Shapiro Wilk tests not satisfied), a Mann Whitney test was used. Multiple comparisons were analyzed using a one-way ANOVA with Bonferroni’s correction (for normal distributions), or a non-parametrical Kruskal-Wallis test followed by a Dunn’s multiple comparisons test. Statistical analyses were conducted with the GraphPad Prism9 software.

Mean are represented and error bars indicate standard deviation (SD) in all figures. The data presented here come from at least three independent experiments. For p values, not significant p>0.05; *p<0.05; **p<0.01; ***p<0.001: ****p<0.0001.

## Supporting information

Supplementary Material Quintin et al. 2022

Movie 01_control_I2_1mM_H2O2

Movie 02_control_PHA_1mM_H2O2

Movie 03_prdx2_I2_1mM_H2O2

Movie 04_prdx2_PHA_1mM_H2O2

Movie 05_control_I2_10microM_H2O2

Movie 06_control_PHA_10microM_H2O2

Movie 07_prdx2_PHA_10microM_H2O2

Movie 08_gur3_I2_1mM_H2O2

Movie 09_gur3_PHA_1mM_H2O2

Movie 10_lite1_I2_1mM_H2O2

Movie 11_lite1_PHA_1mM_H2O2

Movie 12_pmk1_I2_1mM_H2O2

Movie 13_pmk1_PHA_1mM_H2O2

Movie 14_pmk1_I2_10microM_H2O2

Movie 15_pmk1_PHA_10microM_H2O2

Movie 16_sek1_I2_1mM_H2O2

Movie 17_sek1_PHA_1mM_H2O2

Movie 18_sek1_I2_10microM_H2O2

Movie 19_sek1_PHA_10microM_H2O2

Movie 20_I2_control_light_response

Movie 21_PHA_control_light_response

Movie 22_I2&PHA_simulteneous light response

Movie 23_I2_prdx-2mutant_light_response

Movie 24_PHA_prdx-2mutant_light_response

## Acknowledgements

We are grateful to all the staff members of the Imaging Center of the IGBMC, especially Elvire Guiot, Marine Silvin, Erwan Grandgirard and Bertrand Vernay for assistance in confocal microscopy. We thank Christelle Gally, Basile Jacquel and Eric Marois for helpful discussions and critical reading of the manuscript, Sandra Bour for assistance with figure design, Doulaye Dembele for help with statistic analyses. We are indebted to the Reymann, Vermot and Jarriault labs for sharing their equipments and reagents, and for scientific input. We thank the Horvitz lab, especially Na An, for providing the MT GCaMP strains and the Pujol lab for providing the *sek-1* mutant. We thank WormBase for providing resources and the *Caenorhabditis* Genetics Center (funded by NIH Office of Research Infrastructure Programs P40 OD010440, University of Minnesota) for providing strains.

## Author contributions

S.Q. designed and conducted the experiments, analyzed data and wrote the manuscript; T.A. designed and printed the microfluidic chip; T. Y. contributed to ChIP seq data analyses; G.C. developed the Matlab pipeline for data analyses.

