## Supplementary Material Quintin et al. 2022 for "Distinct mechanisms underlie H_2_O_2_ sensing in *C. elegans* head and tail"

#### List of supplementary information

#### Supplementary Figures

**Supplementary Figure 1**, related to Fig 1- The available PRDX-2::GFP transgenic line (CTD1051.3) does not suit to our study.
**Supplementary Figure 2**, related to Figs 1-2- PRDX-2 induction is not observed in I2 neurons upon $H_2O_2$ treatment.
**Supplementary Figure 3**, related to Fig 2- Genomic organization of the *prdx-2* locus and presumptive regulation by SKN-1 and additional transcription factors.
**Supplementary Figure 4**, related to Figs 3-4- Design of the microfluidic chip used in $H_2O_2$ neuronal response experiments.
**Supplementary Figure 5,** related to Fig 3- Individual intensity measurements of movies 1-7.
**Supplementary Figure 6**, related to Fig 3- *prdx-2* mutants PHA neurons do not respond to 10µM $H_2O_2$.
**Supplementary Figure 7**, related to Fig 4- *gur-3* and *lite-1* mutants show reciprocal phenotypes in $H_2O_2$ sensing in I2 and in PHA neurons.
**Supplementary Figure 8,** related to Fig 4- Individual intensity measurements of movies 8-11 (*gur-3* and *lite-1* mutants).
**Supplementary Figure 9,** related to Fig 4, continued - Individual intensity measurements of movies 12-15 (*pmk-1* mutants).
**Supplementary Figure 10,** related to Fig 4, continued - Individual intensity measurements of movies 16-19 (*sek-1* mutants).
**Supplementary Figure 11,** related to Fig 4. Unlike in I2 neurons, PRDX-2 is not essential for light sensing in PHA neurons.
**Supplementary Figure 12,** related to Discussion- A presumptive model of how GUR-3 and LITE-1 receptors may be activated intracellularly by PRDX-2.
**Supplementary Figure 13,** related to Discussion- PHA and ASH neurons belong to the same neuronal cluster, and share partially overlapping transcriptomic signatures.

#### Supplementary Movies

**Movie 1**- wild-type control I2 response to 1mM $H_2O_2$
**Movie 2**- wild-type control PHA response to 1mM $H_2O_2$
**Movie 3**- *prdx-2* mutant I2 response to 1mM $H_2O_2$
**Movie 4**- *prdx-2* mutant PHA response to 1mM $H_2O_2$
**Movie 5**- wild-type control I2 response to 10µM $H_2O_2$
**Movie 6**- wild-type control PHA response to 10µM $H_2O_2$
**Movie 7**- *prdx-2* mutant PHA response to 10µM $H_2O_2$
**Movie 8**- *gur-3* mutant I2 response to 1mM $H_2O_2$
**Movie 9**- *gur-3* mutant PHA response to 1mM $H_2O_2$
**Movie 10**- *lite-1* mutant I2 response to 1mM $H_2O_2$
**Movie 11**- *lite-1* mutant PHA response to 1mM $H_2O_2$
**Movie 12**- *pmk-1* mutant I2 response to 1mM $H_2O_2$
**Movie 13**- *pmk-1* mutant PHA response to 1mM $H_2O_2$
**Movie 14**- *pmk-1* mutant I2 response to 10µM $H_2O_2$

**Movie 15**- *pmk-1* mutant PHA response to 10µM $H_2O_2$
**Movie 16**- *sek-1* mutant I2 response to 1mM $H_2O_2$
**Movie 17**- *sek-1* mutant PHA response to 1mM $H_2O_2$
**Movie 18**- *sek-1* mutant I2 response to 10µM $H_2O_2$
**Movie 19**- *sek-1* mutant PHA response to 10µM $H_2O_2$
**Movie 20**- wild-type control I2 response to light
**Movie 21**- wild-type control PHA response to light
**Movie 22**- wild-type control I2 and PHA simultaneous responses to light
**Movie 23**- *prdx-2* mutant I2 response to light
**Movie 24**- *prdx-2* mutant PHA response to light

**Movie legends, see page 16**

#### Supplementary Methods and Materials

**I- Matlab tutorial for analysis of *C. elegans* neuronal activation using a calcium sensor, page 17**

**II-Analysis of single cell transcriptomic data in ASH and PHA neurons (for the generation of the dot plots in S13 Fig), page 20**

**III- List of strains used (page 21)**

**IV- List of the oligonucleotides used to generate CRISPR/Cas 9 knock-in lines (page 22)**

#### Supplementary Tables (see separate Excel files)

**S1 Table - Differential analysis of genes expressed in cluster 28 versus all neurons**

**S2 Table - Differential analysis of genes expressed in cluster 28.0 (PHA) versus all neurons**

**S3 Table - Differential analysis of genes expressed in cluster 28.1 (ASH) versus all neurons**

#### Supplementary References (page 23)

### Supplementary Figures

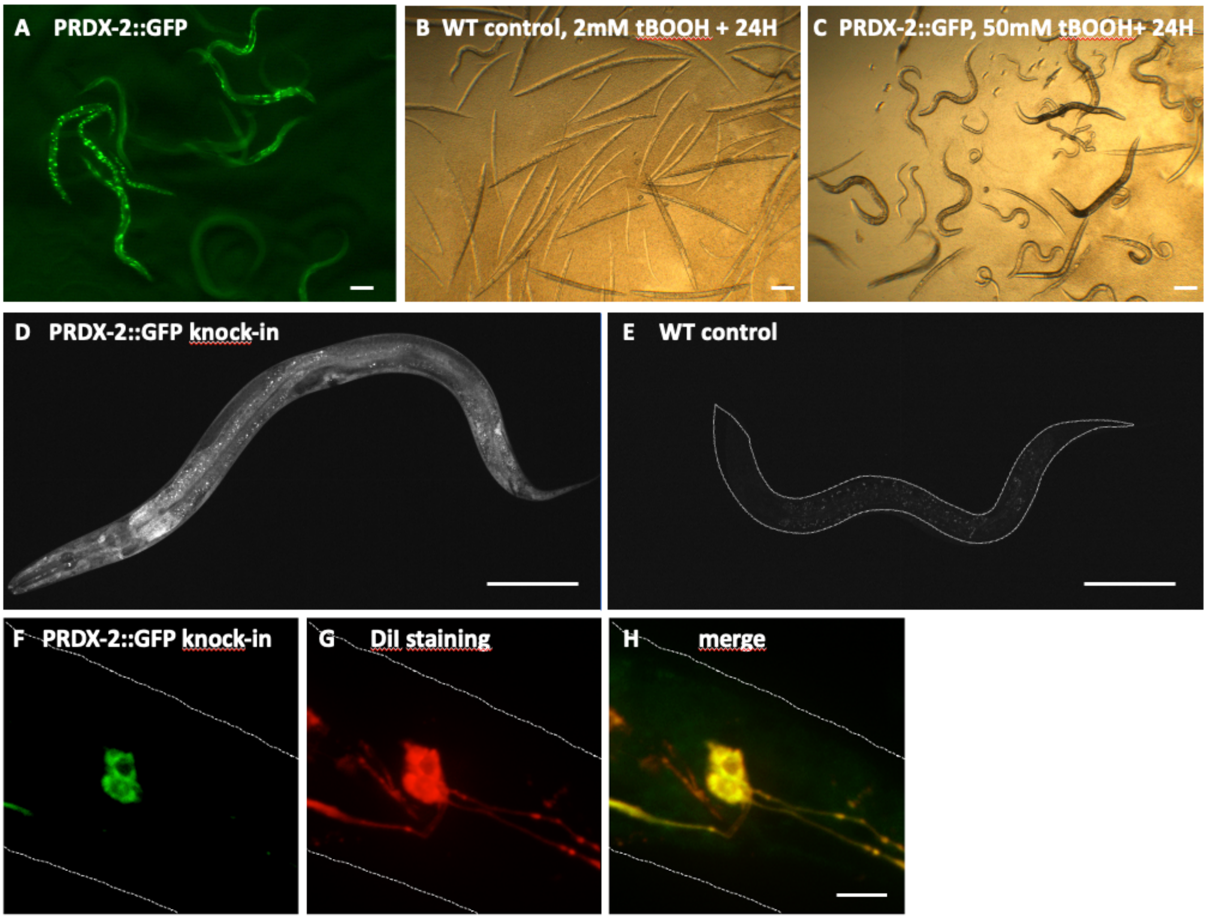


**Supplementary Figure 1, related to Fig 1- The available PRDX-2::GFP transgenic line (CTD1051.3) does not suit to our study.**
(A) Low magnification fluorescent image of untreated animals from the CTD1051.3 line showing that transgenics express aggregates of PRDX-2::GFP, a hallmark of overexpression of the fusion protein. (B,C) Low magnification images of animals treated in flat bottom wells imaged after 24h of treatment in the potent oxidative stress inducing agent tBOOH. In the wild-type control well (A), almost all animals are all dead in 2mM tBOOH, appearing as rods, while CTD1051.3 transgenics survive a 25X higher dose of the drug, indicating their much stronger resistance to oxidative stress. (D) Representative image of an animal of the PRDX-2::GFP knock-in line; a wild-type control imaged with the same settings is shown (E), delineated by a dotted line. Bar, 100µm. (F-H) Confocal projections showing the tail region (dotted contours) of a PRDX-2::GFP transgenic animal stained with the lipophilic orange-red dye DiI (see https://www.wormatlas.org/EMmethods/DiIDiO.htm), imaged in green (F) and red channels (G). The overlay (H) shows that PRDX-2::GFP tail neurons are stained by the dye, establishing their identity as phasmid sensory neurons (PHA/PHB). Bar, 10µm.

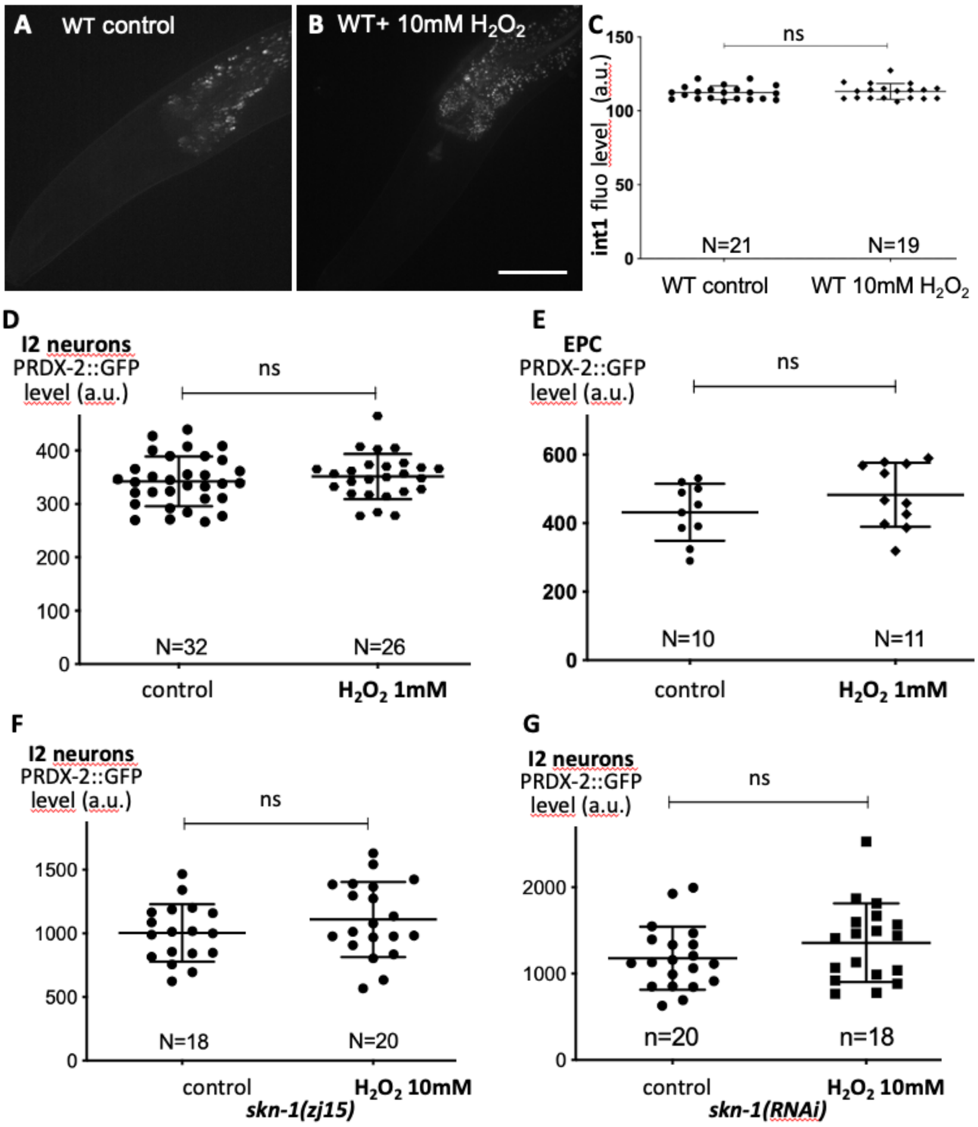


**Supplementary Figure 2, related to Figs 1, 2- PRDX-2 induction is not observed in I2 neurons upon $H_2O_2$ treatment.**
(A,B) Wild-type controls do not show a higher gut autofluorescence after $H_2O_2$ treatment, as illustrated by the int1 anterior gut cell fluorescence quantification (C). (D,E) Quantification of the PRDX-2::GFP fluorescence level in I2 neurons and in the excretory pore cell (EPC) in controls after 1mM-$H_2O_2$ treatment. (F,G) Quantification of I2 neurons' PRDX-2::GFP fluorescence level in *skn-1(zj15)* mutants and in *skn-1(RNAi)* animals upon a 10mM-$H_2O_2$ treatment. Means are shown and error bars represent SD; ns, not significant, $p>0.05$ (t test or Mann-Whitney test). Scale bar, 50µm.

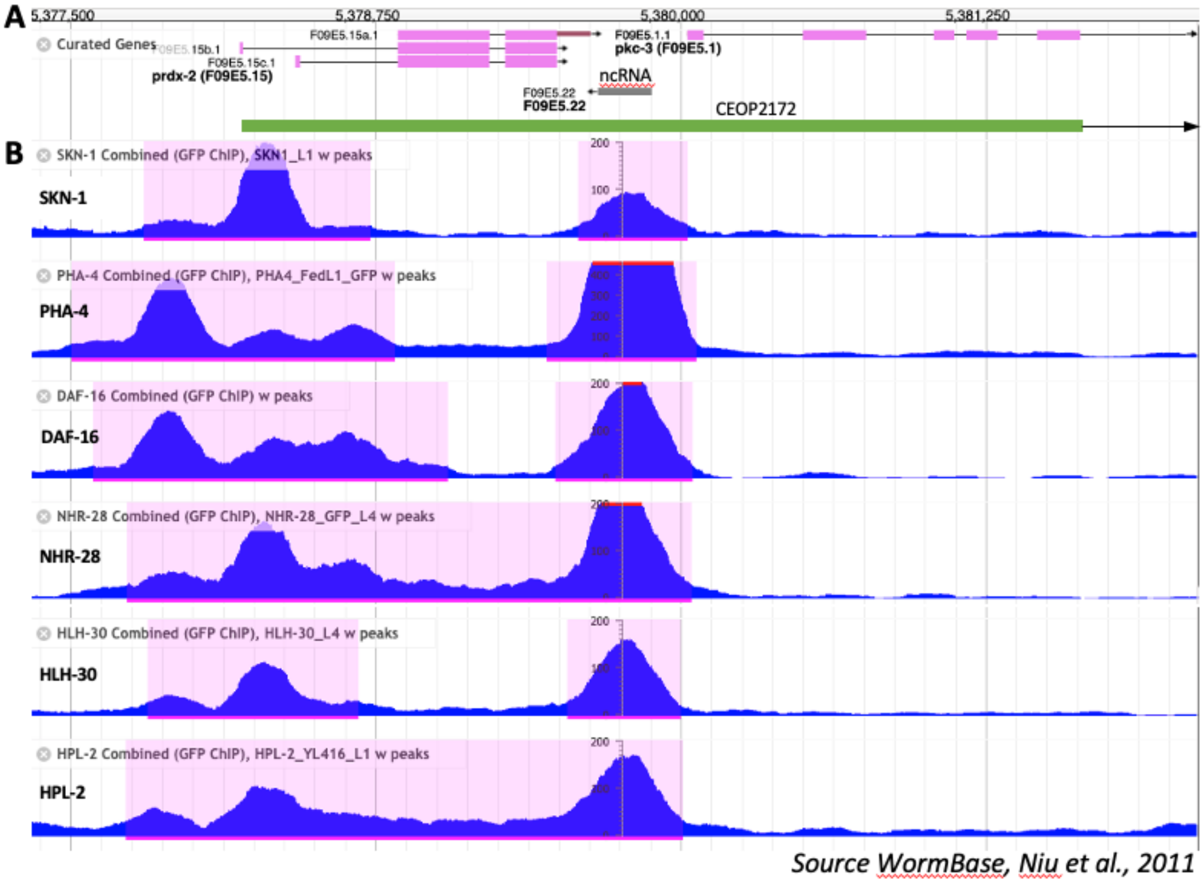


**Supplemental Figure 3, related to Fig 2. Genomic organization of the *prdx-2* locus and presumptive regulation by SKN-1 and additional transcription factors.** (A) Genome Browser screenshot from WormBase (release WS283, JBrowse II), showing the gene organization on chromosome II, at the indicated coordinates (top), and the peaks detected by ChIP-seq using an anti-GFP antibody (Niu et al. 2011) in *prdx-2* and *pkc-3* promoters, in GFP-tagged transgenic lines of the indicated transcription factors (B). Note the co-regulation of *prdx-2* and *pkc-3*, which are organized in an operon, illustrated by the green bar.

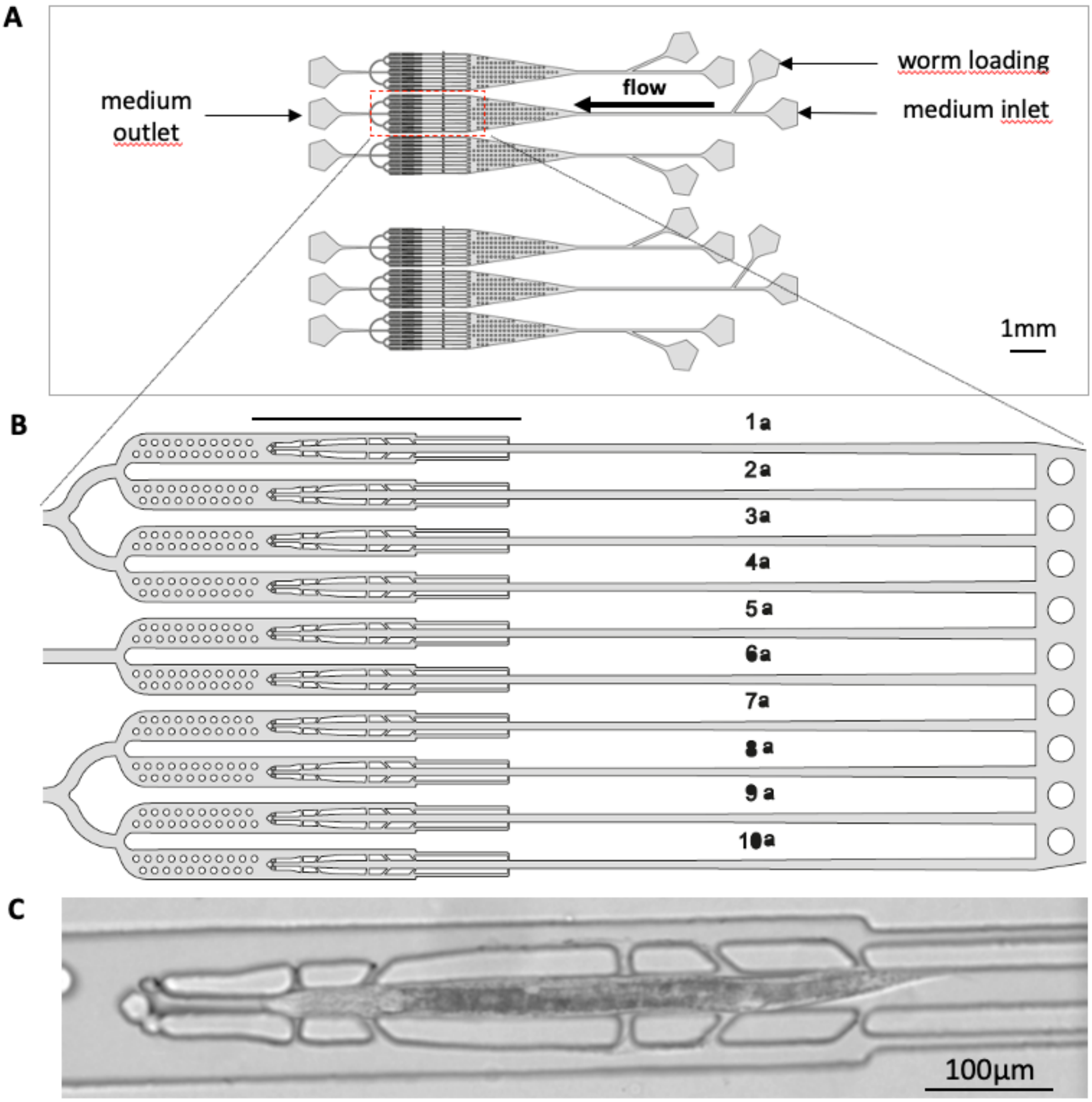


**Supplementary Figure 4, related to Figs 3-4- Design of the microfluidic chip used in $H_2O_2$ neuron response experiments.**

(A) Global view of the microfluidic chip used in all neuron response experiments, showing the 6 independent series of 10 worm traps. The original file (Autocad format) is available at https://github.com/gcharvin/viewworm.

(B) Magnification of the worm trap area showing the 10 individual channels (corresponding to the red box in A). Scale bar, 1mm

(C) DIC image of an animal trapped (anterior to the left). Occasionally animals were trapped in the opposite orientation, but this did not affect the neuronal response. Scale bar, 100µm.

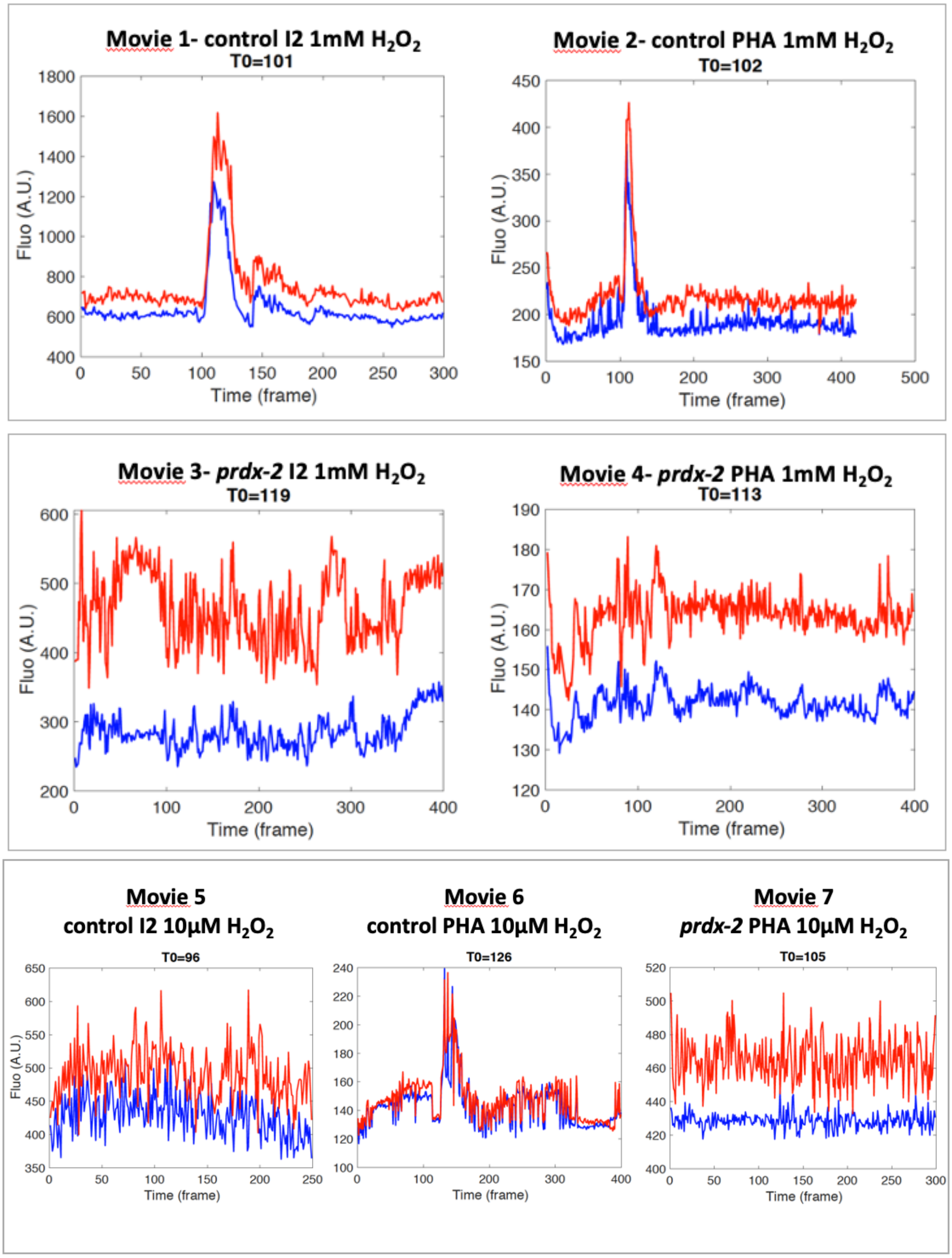


**Supplementary Figure 5, related to Fig 3- Individual intensity measurements of Movies 1-7.**

The curves represent the mean GCaMP3 intensity raw value over time (1 frame=2sec) quantified in I2 and PHA neurons in control and *prdx-2* mutants corresponding movies (1-7), upon a 1mM or a 10µM $H_2O_2$ exposure. T0 indicate the time point at which the $H_2O_2$ treatment has been applied during 100 frames. Red and blue colors represent left and right I2 and PHA neurons (*ie* I2L/R and PHAL/R). Note that the colors have been changed in normalized average curves shown in Fig 3 (Movies 1-6) and in related S6 Fig (Movie 7).

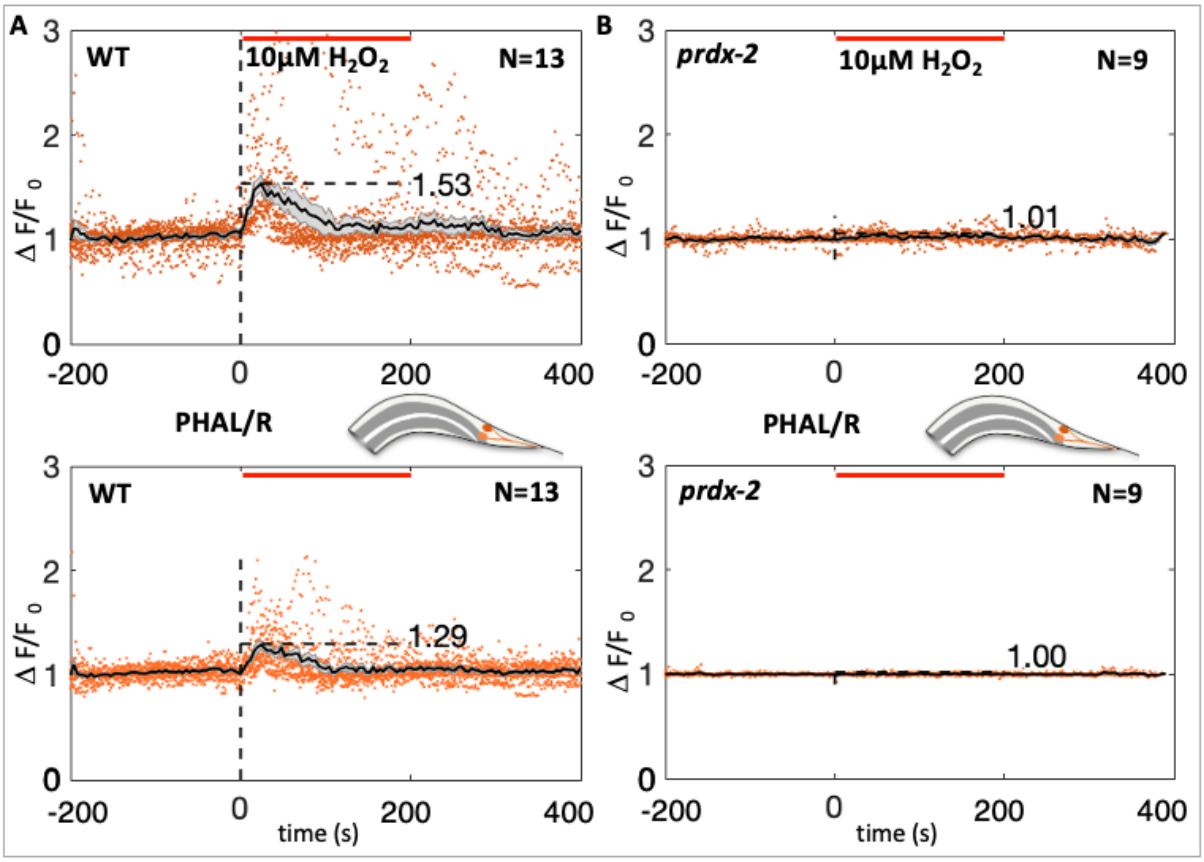


**Supplementary Figure 6, related to Fig 3- *prdx-2* mutants PHA neurons do not respond to 10µM $H_2O_2$.**

(A-B) Average curves showing the normalized calcium response to 10µM $H_2O_2$ measured over time (in seconds) using the GCaMP3 sensor in PHA left and right neurons (top and bottom curves) in wild-type controls (A) and in *prdx-2(gk169)* mutants (B). N, number of movies analyzed for each genotype. See Movies 6-7 and related S5 Fig.

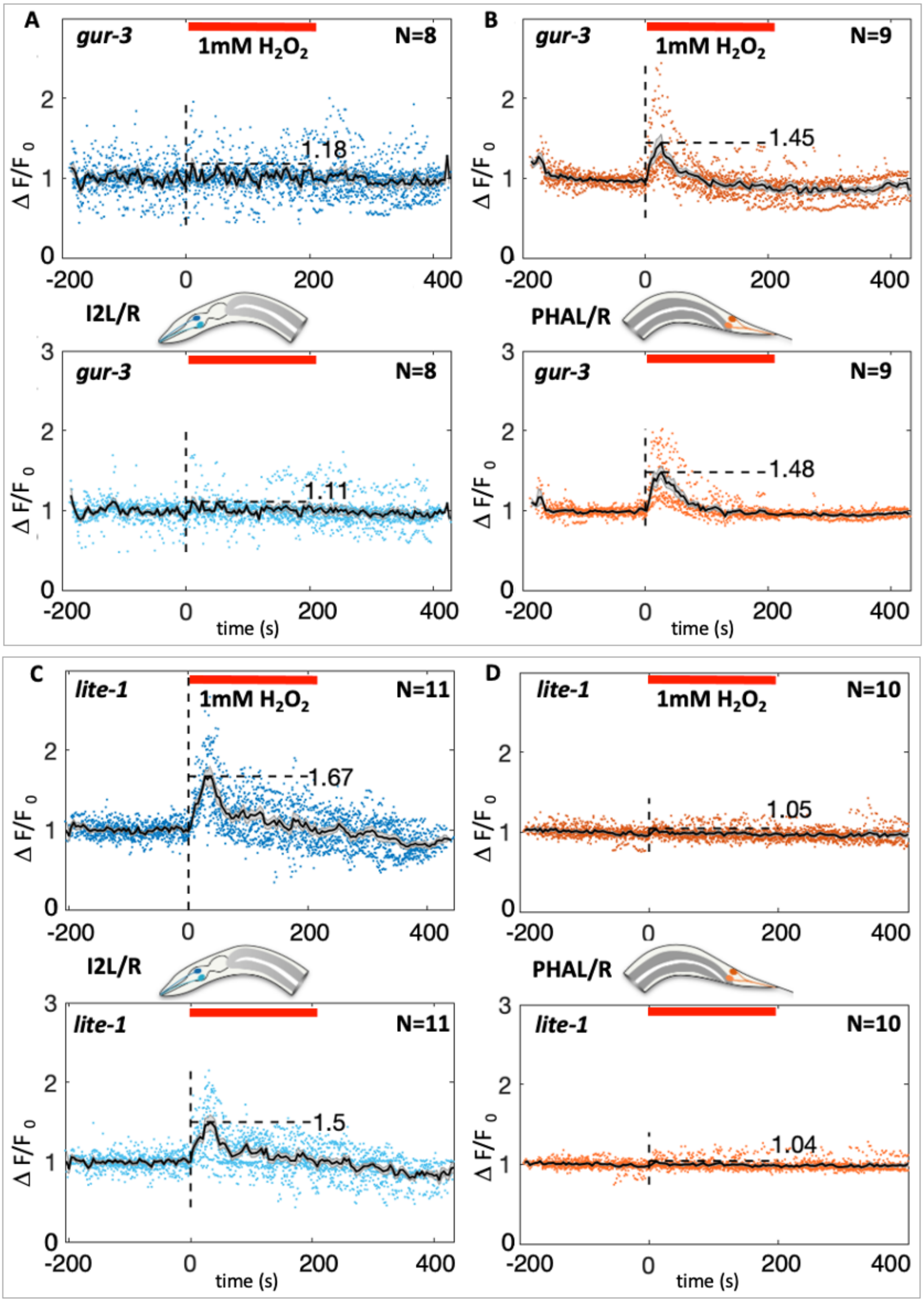


**Supplementary Figure 7, related to Fig 4- *gur-3* and *lite-1* mutants show reciprocal phenotypes in $H_2O_2$ sensing in I2 and in PHA neurons.**
(A-D) Average curves showing the normalized calcium response to 1mM $H_2O_2$ measured over time (indicated in seconds) using the GCaMP3 sensor in I2 and PHA left and right neurons (top and bottom curves) in *gur-3(ok2245)* (A,B) and *lite-1(ce314)* mutants (C,D). N, number of movies analyzed for each genotype. See Movies 8-11 and related S8 Fig.

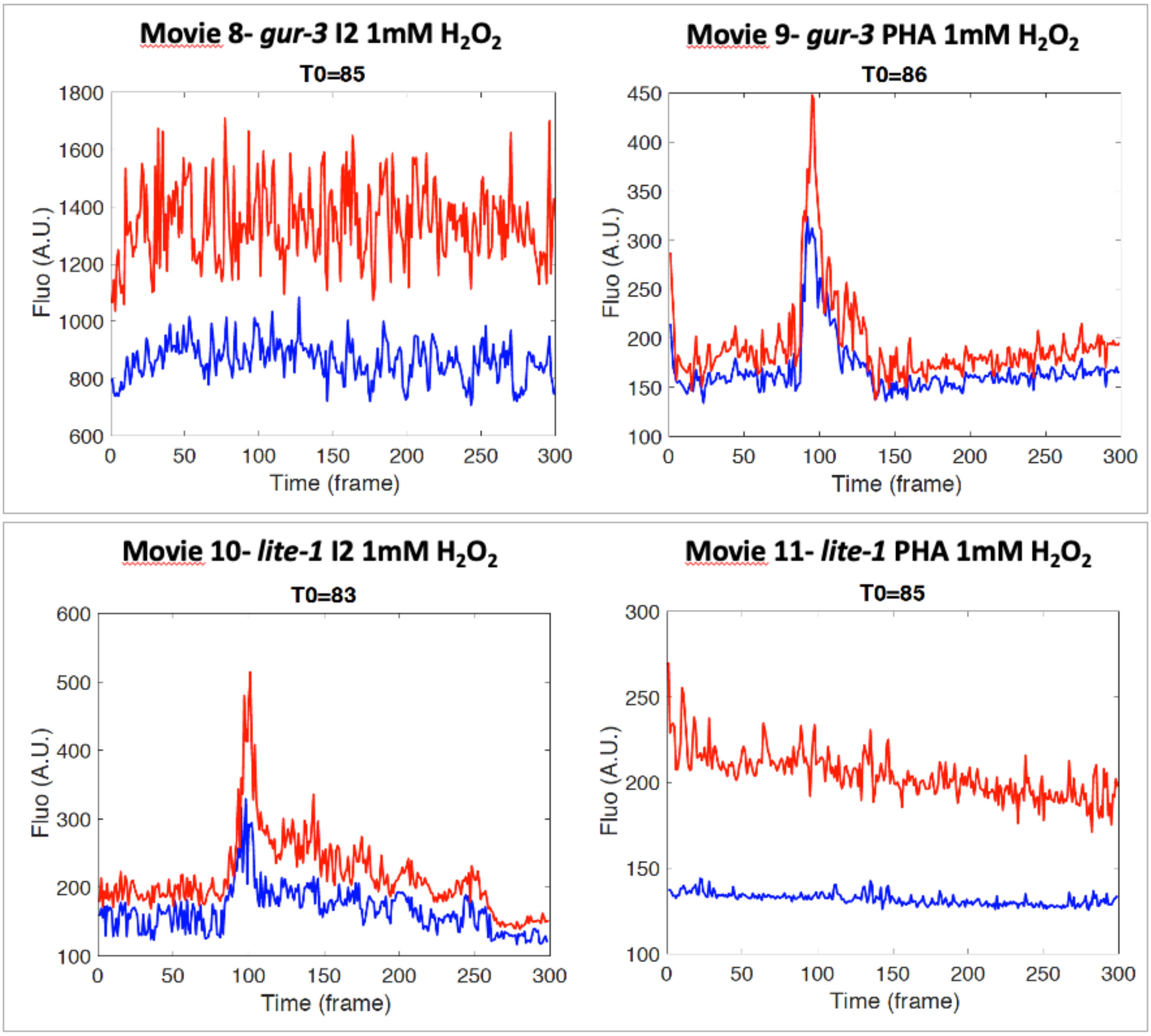


**Supplementary Figure 8, related to Fig 4- Individual intensity measurements of Movies 8-11 (*gur-3* and *lite-1* mutants).**

The curves represent the mean GCaMP3 intensity raw value over time (1 frame=2sec) quantified in I2 and PHA neurons in *gur-3* and *lite-1* mutants upon a 1mM $H_2O_2$ exposure (starting at T0 and lasting 100 frames), in corresponding movies (8-11). Red and blue colors show left and right neurons for I2 (left panel) and PHA (right panel) neurons (*ie* I2L/R and PHAL/R). Note the reciprocal phenotypes observed in both mutants. Colors have been changed in the related normalized average curves shown in S7 Fig.

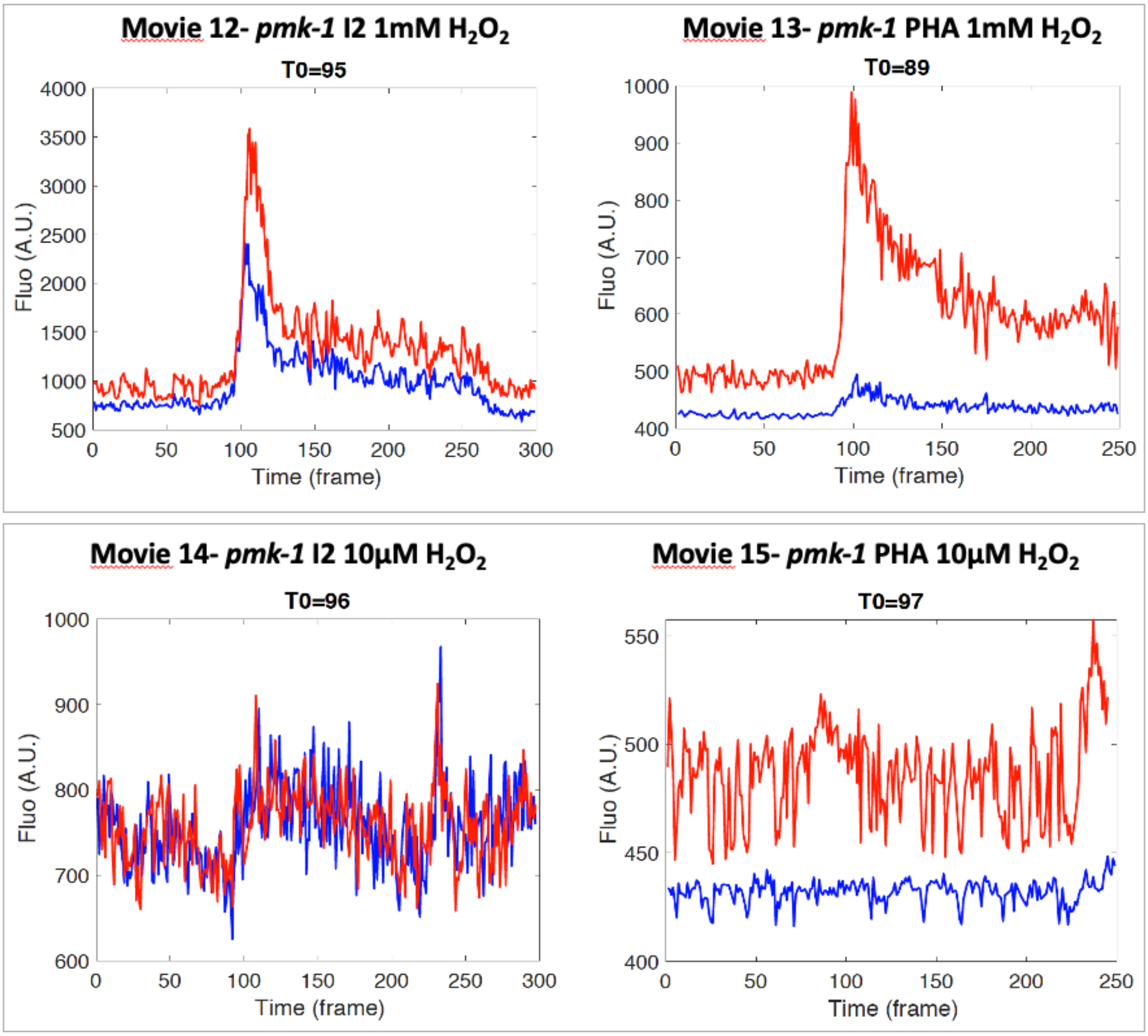


**Supplementary Figure 9, related to Fig 4, continued- Individual intensity measurements of Movies 12-15 (*pmk-1* mutants).**

The curves represent the mean GCaMP3 intensity raw value over time (1 frame=2sec) quantified in I2 and PHA neurons in *pmk-1* mutants upon a 1mM or a 10µM $H_2O_2$ stimulation (starting at T0 and lasting 100 frames), in corresponding movies (12-15). Red and blue color indicate I2L/R neurons (left panel) and PHAL/R neurons (right panel).

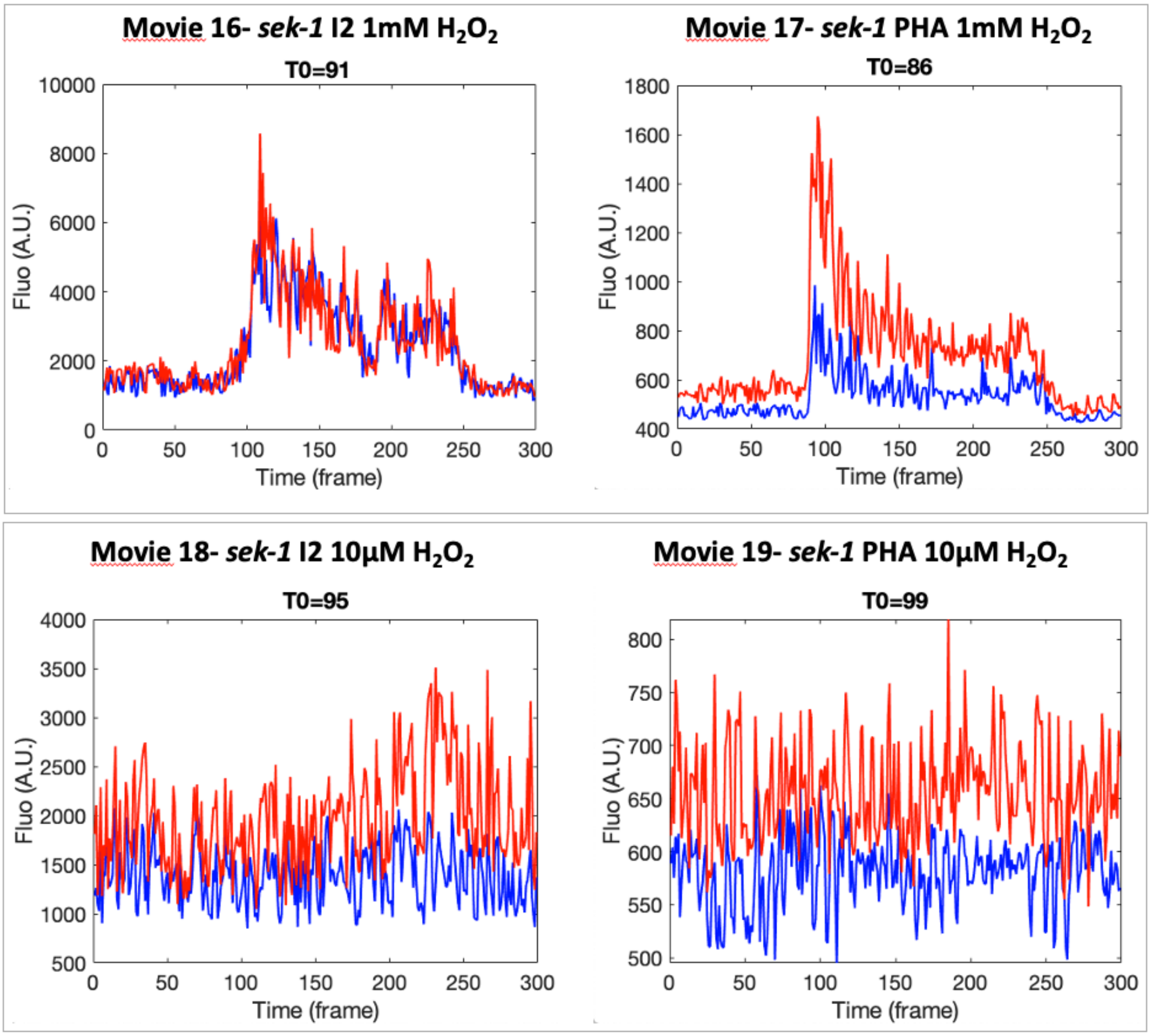


**Supplementary Figure 10, related to Fig 4, continued- Individual intensity measurements of Movies 16-19 (*sek-1* mutants).**
The curves represent the mean GCaMP3 intensity raw value over time (1 frame=2sec) quantified in I2 and PHA neurons in *sek-1* mutants upon a 1mM or a 10µM $H_2O_2$ stimulation (starting at T0 and lasting 100 frames), in corresponding movies (16-19). Red and blue color indicate left and right I2 (left panel) and PHA (right panel) neurons.

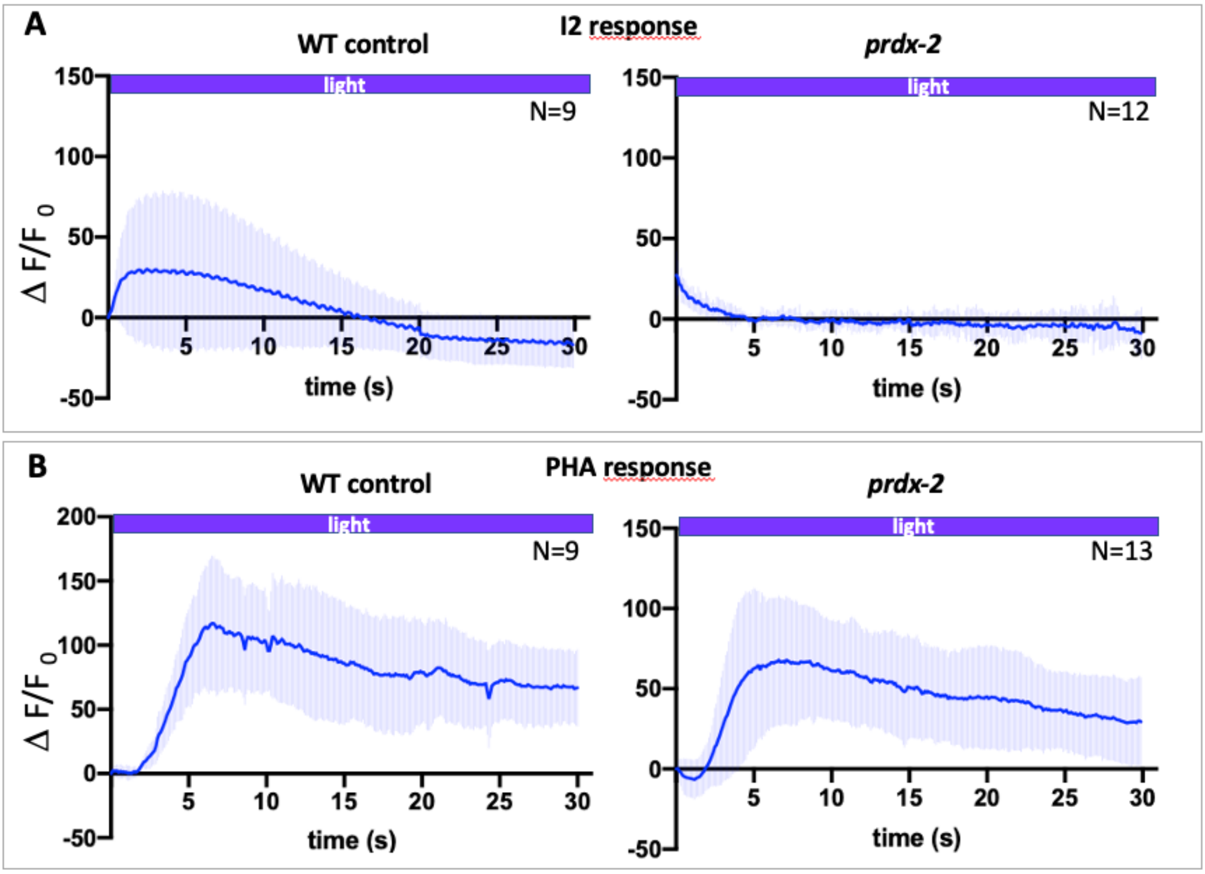


**Supplementary Figure 11, related to Fig. 4, continued - Unlike in I2 neurons, PRDX-2 is not essential for light sensing in PHA neurons.**
Average curves showing the normalized calcium response to blue light over time (in seconds) using the GCaMP3 sensor, measured in the soma of I2 (A) and PHA neurons (B) in wild-type controls and in *prdx-2(gk169)* mutants. N, number of movies analyzed for each genotype. Error bars represent SD. While I2 neurons fail to respond to light in *prdx-2* mutants, PHA neurons do respond, albeit with a lower intensity peak than in controls. See related Movies 20-24.

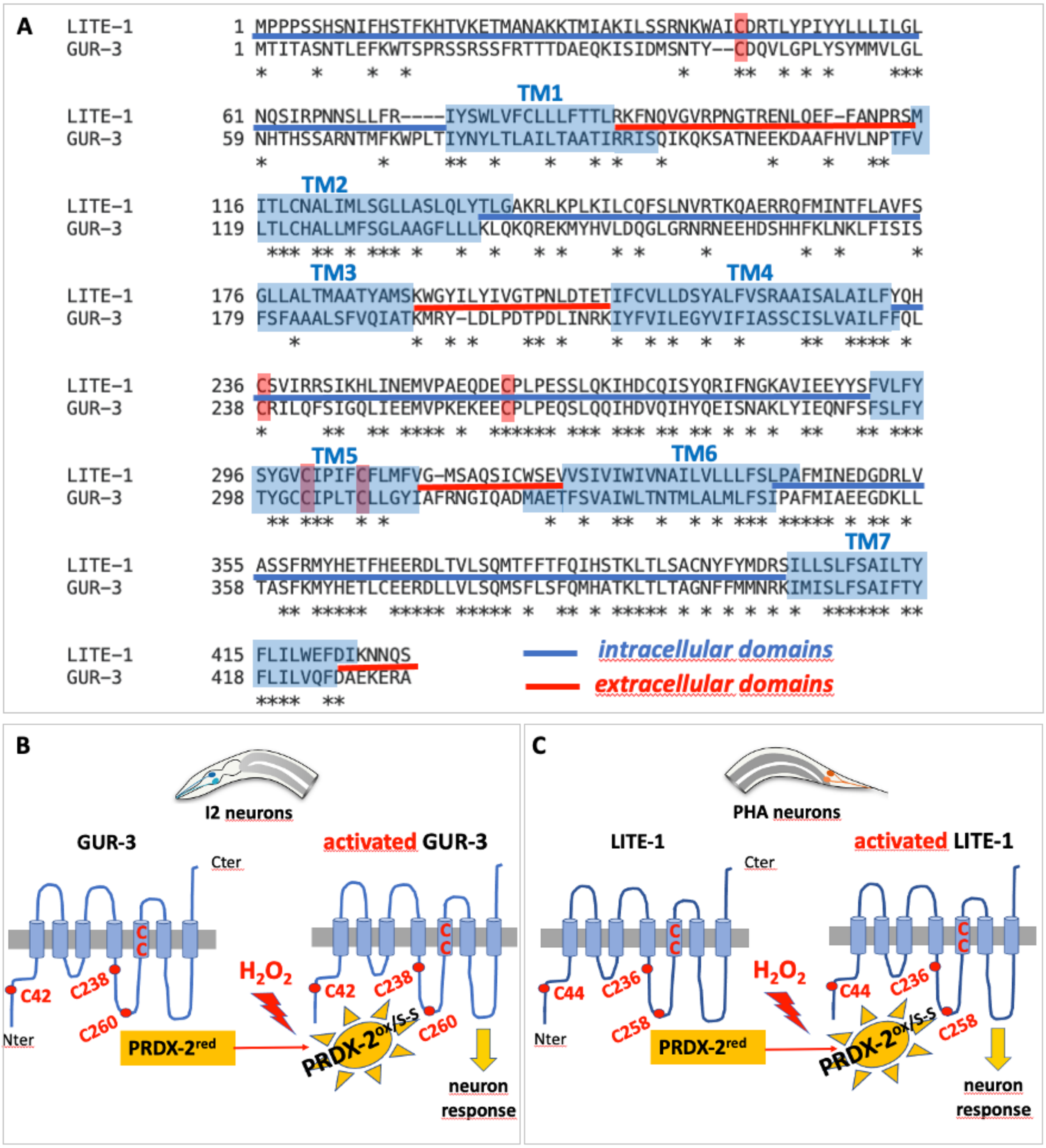


**Supplementary Figure 12, related to Discussion- A presumptive model of how GUR-3 and LITE-1 receptors may be activated intracellularly by PRDX-2.**
(A) Alignment of LITE-1 and GUR-3 protein sequences made with the SIM alignment tool (https://web.expasy.org/sim/), using the comparison matrix BLOSUM30. Transmembrane (TM) and intra/extracellular domains were predicted using the DeepTMHMM program (https://dtu.biolib.com/DeepTMHMM). The alignment reveals 39.9% identity over 434 residues overlap. Note the rather large intracellular domains (in blue), encompassing conserved cysteines (boxed in red) (B-C) A putative PRDX-2 redox relay may trigger $H_2O_2$-induced receptor activation in I2 and PHA neurons. Sketch of GUR-3 (B) and LITE-1 (C) receptors, deduced from A, depicting their conserved cysteines. Upon $H_2O_2$ exposure, oxidized PRDX-2 or its disulfide form (PRDX-$2^{ox/S-S}$) could oxidize these cysteines, possibly forming a disulfide conjugate and/or inducing a conformation change, which would subsequently trigger receptor activation, and I2 or PHA neuron response.

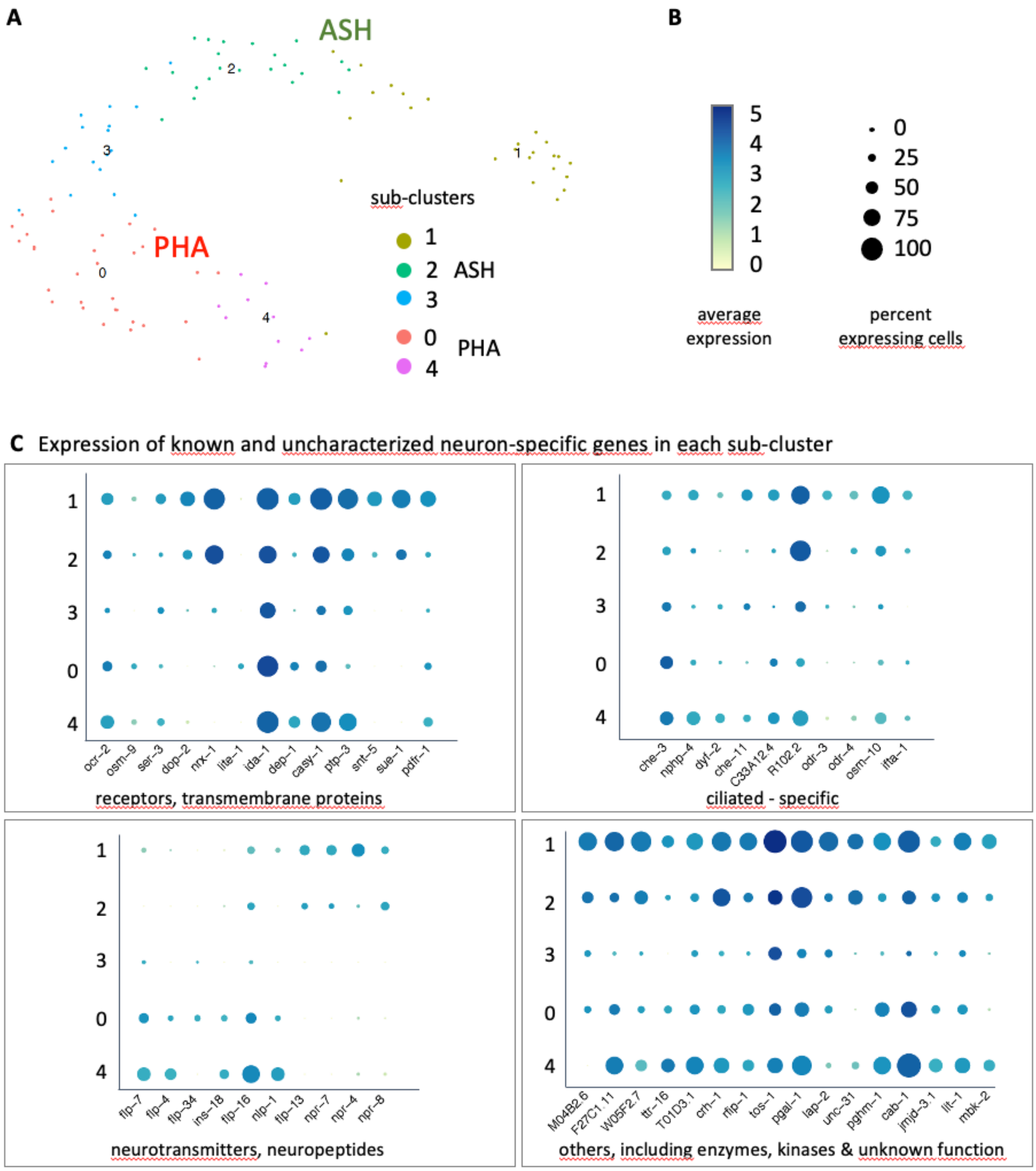


**Supplementary Figure 13, related to Discussion- PHA and ASH neurons belong to the same neuronal cluster, and share partially overlapping transcriptomic signatures.**
(A)- Uniform Manifold Approximation and Projection (UMAP) projection of parent cluster 28, from (Lorenzo et al. 2020), in which ASH and PHA/PHB nociceptive neurons were found in distinct sub-clusters (Louvain clustering at 8 PCs and at 1.2 resolution).
(B, C)- Dot plot indicating for a selection of genes both the intensity of gene expression and the fraction of expressing cells in each sub-cluster (B), based on single-cell RNA-sequencing data from (Cao et al. 2017). See Supplementary Methods, p. 20 and related S1-S3 Tables for exhaustive lists of genes expressed in each cluster.

#### Movie Legends

**Movie 1**- wild-type control I2 response to 1mM $H_2O_2$
**Movie 2**- wild-type control PHA response to 1mM $H_2O_2$
**Movie 3**- *prdx-2* mutant I2 response to 1mM $H_2O_2$
**Movie 4**- *prdx-2* mutant PHA response to 1mM $H_2O_2$
**Movie 5**- wild-type control I2 response to 10µM $H_2O_2$
**Movie 6**- wild-type control PHA response to 10µM $H_2O_2$
**Movie 7**- *prdx-2* mutant PHA response to 10µM $H_2O_2$
**Movie 8**- *gur-3* mutant I2 response to 1mM $H_2O_2$
**Movie 9**- *gur-3* mutant PHA response to 1mM $H_2O_2$
**Movie 10**- *lite-1* mutant I2 response to 1mM $H_2O_2$
**Movie 11**- *lite-1* mutant PHA response to 1mM $H_2O_2$
**Movie 12**- *pmk-1* mutant I2 response to 1mM $H_2O_2$
**Movie 13**- *pmk-1* mutant PHA response to 1mM $H_2O_2$
**Movie 14**- *pmk-1* mutant I2 response to 10µM $H_2O_2$
**Movie 15**- *pmk-1* mutant PHA response to 10µM $H_2O_2$
**Movie 16**- *sek-1* mutant I2 response to 1mM $H_2O_2$
**Movie 17**- *sek-1* mutant PHA response to 1mM $H_2O_2$
**Movie 18**- *sek-1* mutant I2 response to 10µM $H_2O_2$
**Movie 19**- *sek-1* mutant PHA response to 10µM $H_2O_2$

**Legend of Movies 1-19 (related to Figs 3, 4 and S5, S8-S10 Figs)**
All movies show the neuronal responses to $H_2O_2$ (at the indicated dose) in I2 and PHA neurons visualized using the GCaMP3 calcium sensor, in animals trapped in microfluidic chambers. All movies shown have been processed for re-alignment to facilitate response visualization (using the Matlab Readworm_PHA code), and accelerated 60 times. See corresponding fluorescence intensity measurements of each of these movies in S5 and S8-S10 Figs.

**Movie 20**- wild-type control I2 response to light
**Movie 21**- wild-type control PHA response to light
**Movie 22**- wild-type control I2 and PHA simultaneous responses to light
**Legend of Movies 20-22 (related to Fig 4)**
Neuronal responses to blue light (485nm) in I2 and PHA neurons visualized with the GCaMP3 calcium sensor in wild-type control animals. Original movies have been colored artificially to highlight fluorescence intensity variations (using Fiji 'glow' look up table). Note the different pace of response of each neuron, especially in the simultaneous recording (Movie 22), and the contrast between the quasi instantaneous response of the posterior neurite in I2 neurons (movies 20,22, Fig 4G) and the strikingly long soma response in PHA (Movies 21,22, Fig 4H). Accelerated 5 times.

**Movie 23**- *prdx-2* mutant I2 response to light
**Movie 24**- *prdx-2* mutant PHA response to light
**Legend of Movies 23-24 (related to S11 Fig)**
Neuronal responses to blue light (485nm) in *prdx-2* mutants I2 and PHA neurons, visualized with the GCaMP3 calcium sensor. Original movies have been colored artificially to highlight fluorescence intensity variations (Fiji 'glow' look up table). Note that I2 neurons show no response to light (movie 23), while PHA neurons respond normally (movie 24) in *prdx-2* mutants. Accelerated 5 times.

#### Supplementary Methods and Materials

**I- Matlab tutorial for analysis of *C. elegans* neuronal activation using a calcium sensor**

These codes have been created for tracking timelapse recordings of left and right neuronal pairs expressing a calcium sensor such as GCaMP, and available at: https://github.com/gcharvin/viewworm. Below are described the sequential functions to run under Matlab to perform movie quantification.

Upon acquisition, movies should be cropped as small as possible to minimize processing time and should be save as .tif files.
**Note**: we recommend to first make a 4D-projection of the original movie, draw the smallest ROI around the neurons and transfer it to the original movie, to ensure that all frames are properly included in the cropped selection.

##### 1. Movie registration

**—> readworm_PHA ('movie_name.tif')**

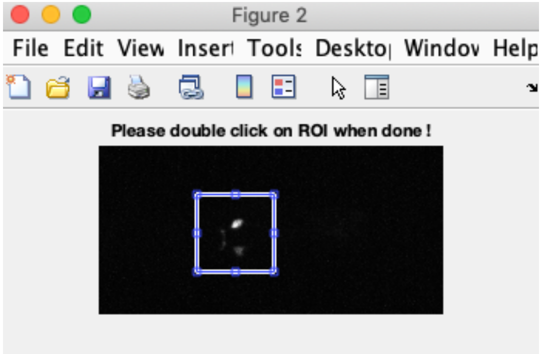


After ROI selection with the 2 neurons of interest (shown above), the function readworm_PHA processes every movie time point and performs the alignment of neurons, generating two files:

movie-registered.mat
movie.mp4

The mp4 allows the user to quickly verify that the movie has been correctly aligned.
**Note**: supplemental movies 1-14 were generated using this program.

##### 2. Neuron segmentation, selection and tracking

**—> viewworm_2('movie_name')**

The function opens a figure project with several images and buttons.
The top left image allows movie traveling in the registered movie for each frame, using the right and left keyboard arrows, or typing directly the frame number in the box at the bottom of the window. To better visualize neurons, the intensity threshold of the image viewer can be adjusted with the top and bottom keyboard arrows.

Using the '**Pixel Train**' button, the user teaches the program which pixels should be selected for neuronal segmentation by painting them in green (left mouse click). The pixels which should not be included for segmentation are painted in red (right mouse click). This needs to be done for several time points to obtain the best segmentation results, especially during the neuronal response. Below is shown an example with the selected neurons in green and the posterior neurite in red (excluded from the analysis), during the response.

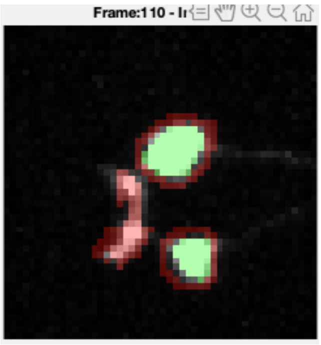


When this teaching has been done as often as needed, the user clicks ‘**Classify pixels**’, leading to the **neuron segmentation** as defined by the user. Then the user defines each neuron in the bottom segmented image, by clicking the ‘**Select neurons**‘ button:

right click—> red selection (left neuron);
left click —> blue neuron (right neuron);

Upon pressing the ‘**Track’** button, left and right neurons will be tracked for all movie time points. Upon pressing ‘**Plot’**, the **mean fluorescence intensity** is indicated as a function of time, generating the red and blue curves in the bottom right panel, as shown below.

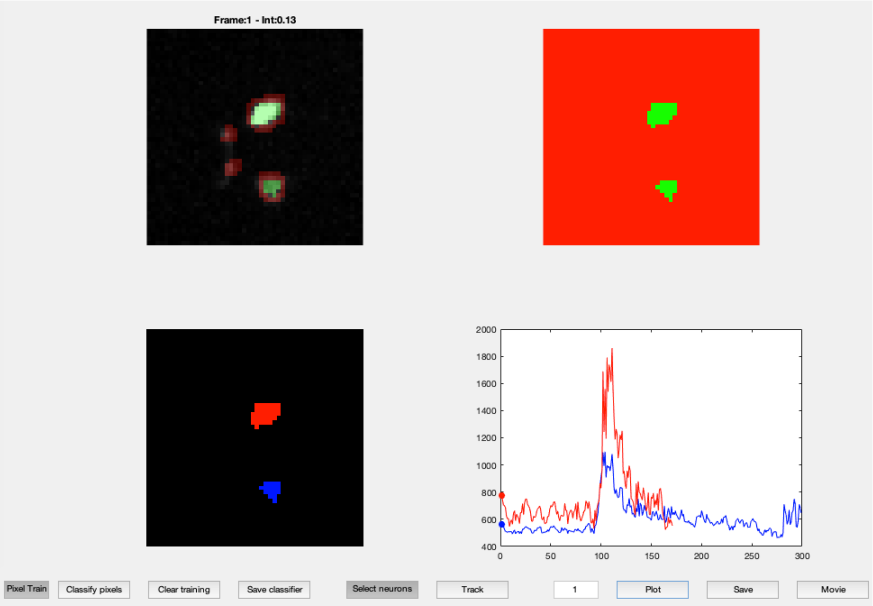


**Important note** : this step allows the **correction of tracking errors** at every frame. The quality control is performed by the user by selecting the appropriate neuron and clicking ‘**Track’** again until all tracking errors are corrected. Below is shown an example of error correction at time point 164 (pixels colored in green are excluded from the quantification in the bottom left box):

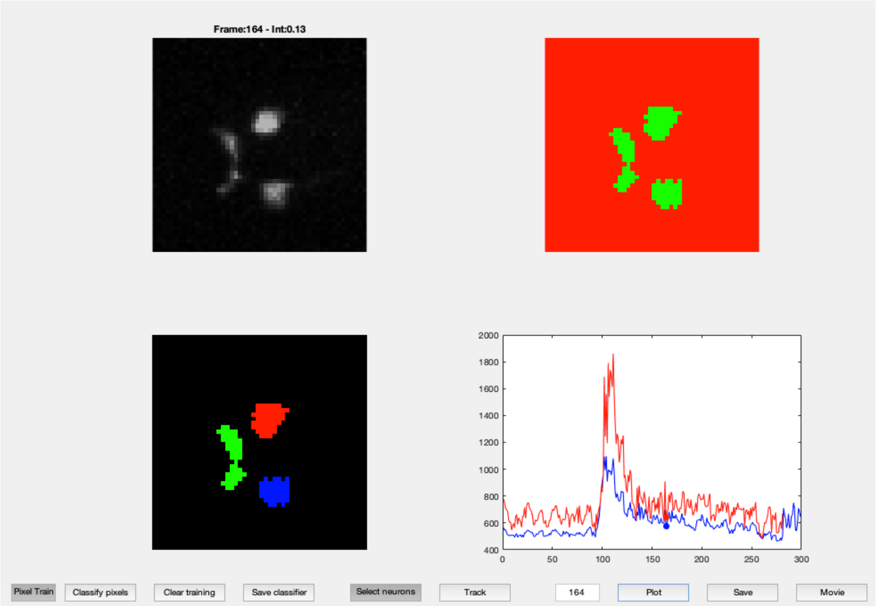


When the movie is properly tracked to the end, click ‘**Save’** button to obtain the .fig files which will be used for subsequent analyses. The user **needs to indicate the T0** in the command window, which corresponds to the time point at which the stress/ stimulus was applied.

Finally click on ’**Movie’** to generate an mp4 movie showing the registered, segmented and tracked movies altogether, allowing an assessment of the global quality of the analysis.

Once all movies have been successfully processed, create a folder containing all the individual .fig analyzed files, which will be used for processing the average curve.

#### 3. Average curve processing

**—> averageGCamp2(‘folder_name')**

In the command window, assign [low, high, offset]=averageGCamp2(‘folder_name’)

**—> processAverage(‘low,high,offset')**

This function will generate average curves for red and blue neurons, for raw and normalized data.

Finally to retrieve individual intensity mean values at the response peak, run the **ExportSingleWormData** function which will produce an Excel sheet with all individual peak values.

#### II- Analysis of single cell transcriptomic data in ASH and PHA neurons (for the generation of the dot plots in S13 Fig)

Single-cell RNA-sequencing data was generated by (Cao et al. 2017). Clusters and sub-clusters were defined as described in (Lorenzo et al. 2020), using Louvain clustering at 8 PCs and at 1.2 resolution. The dataset normalization was performed using Seurat v4 (Hao et al. 2021), and visualized using dot plots. We performed differential analyses using the 'FindMarkers' function of Seurat v4 to compare cluster 28 versus all neurons (data in S1 Table); cluster 28.0 (PHA) versus all neurons (data in S2 Table); clusters 28.1 (ASH) versus all neurons (data in S3 Table). Selected genes were chosen among the most significantly enriched from these lists.

**III- List of strains used**

**N2** wild type, reference strain
**CTD1051.3** N2 *whEx127* [fNH059 (PRDX-2::GFP)+pRF4 *(rol-6(su1006)*] (Hirani et al. 2013)
**SXB01** *prdx-2*(*gch01*[PRDX-2::GFP, *unc-119*(+)]) II; *unc-119(ed3)* III
**SXB05** *prdx-2*(*gch03*[PRDX-2::GFP, *unc-119*(+)] II; *unc-119(ed3)* III
**SXB15** 5 times outcrossed SXB01 *prdx-2*(*gch01*[PRDX-2::GFP, *unc-119*(+)] II
**SXB19** 5 times outcrossed SXB05 *prdx-2*(*gch03*[PRDX-2::GFP, *unc-119*(+)] II
**HT1593** 7 times outcrossed *unc-119(ed3)* III (Dickinson et al. 2013)
**QV225** *skn-1(zj15)* IV (Tang, Dodd, and Choe 2016)
**SXB21** *prdx-2(gch03)* II; *skn-1(zj15)* IV
**MT21650** *nIs575*[*flp-15*prom::GCaMP3, *lin-15*(+)] III; *lin-15(n765)* X (Bhatla and Horvitz 2015)
**MT21570** *nIs575* III; *lite-1(ce314) lin-15(n765)* X (Bhatla and Horvitz 2015)
**MT21785** *nIs575* III; *gur-3(ok2245) lin-15(n765)* X (Bhatla and Horvitz 2015)
**KU4** *sek-1(km4)* IV (Kim et al. 2002)
**KU25** *pmk-1(km25)* IV (Mizuno et al. 2004)
**VC289** *prdx-2(gk169)* II (The *C. elegans* Deletion Mutant Consortium et al. 2012)
**SXB53** *prdx-2(gk169)* II; *nIs575* III
**SXB63** *unc-119(ed3)* III*; gur-3(gch16*[GUR-3::GFP, *unc-119*(+)]*)* X
**SXB67** *nIs575* III; *pmk-1(km25)* IV
**SXB70** *nIs575* III; *sek-1(km4)* X

All SXB strains were made in the course of this study.

#### IV- List of the oligonucleotides used to generate CRISPR/Cas 9 knock-in lines

All primers listed below are in the 5'-3'orientation.

1°) **PRDX-2::GFP knock-in**

Introduction of *prdx-2* sgRNA (into pMLS256):

**Forward**: **TTG**AGTGCTTCTTGAAGTACTCT

**Reverse**: **AAC**AGAGTACTTCAAGAAGCACT

PCR amplification of 5' and 3' homology arms (HAs) of *prdx-2* (including **SapI restriction** site and template-specific overhangs):

— 5' HA PCR product (542bp), upstream *prdx-2* stop codon:

**Forward**: TGT**GCTCTTC**TTggACCCACTTGACTTCACT

**Reverse**: gtg**GCTCTTC**gCGCGTGCTTCTTAAAGTATTCTTGACTTTCTTTGA (with silent mutations to change the PAM sequence)

— 3' HA PCR product (570bp), starting at *prdx-2* stop codon:

**Forward**: act**GCTCTTC**G**ggt**TAAatgtcttacatctctaatttcc

**Reverse**: tta**GCTCTTC**T**tac**atctccgtcctcctctaatgatgt

2°) **GUR-3::GFP knock-in**

Introduction of *gur-3* sgRNA :

**Forward**: **TTG**taagacaattttatTTACAC

**Reverse**: **AAC**GTGTAAataaaattgtctta

PCR amplification of 5' and 3' HAs of *gur-3* (with **SapI restriction** site and template-specific overhangs):

— 5' HA (614bp), upstream *gur-3* stop codon:

**Forward**: tga**GCTCTTC**a**TGG**gttatcatttaaccgtgattgcatgca

**Reverse**: gac**GCTCTTC**t**CGC**CACAGGTGGTTGGACAATGAGCA

— 3' HA (555bp), downstream *gur-3* stop codon:

**Forward**: tcc**GCTCTTC**t**GGT**TAAataaaattgtcttaacattttccccatattga

**Reverse**: ttt**GCTCTTC**g**TAC**tctcgtttaacgatttctcctgtctga

Hirani, Nisha, Marcel Westenberg, Minaxi S. Gami, Paul Davis, Ian A. Hope, and Colin T. Dolphin. 2013. "A Simplified Counter-Selection Recombineering Protocol for Creating Fluorescent Protein Reporter Constructs Directly from *C. Elegans* Fosmid Genomic Clones." *BMC Biotechnology* 13 (January): 1.

Kim, Dennis H., Rhonda Feinbaum, Geneviève Alloing, Fred E. Emerson, Danielle A. Garsin, Hideki Inoue, Miho Tanaka-Hino, et al. 2002. "A Conserved P38 MAP Kinase Pathway in *Caenorhabditis Elegans* Innate Immunity." *Science* 297 (5581): 623–26.

Lorenzo, Ramiro, Michiho Onizuka, Matthieu Defrance, and Patrick Laurent. 2020. "Combining Single-Cell RNA-Sequencing with a Molecular Atlas Unveils New Markers for *Caenorhabditis Elegans* Neuron Classes." *Nucleic Acids Research* 48 (13): 7119–34.

Mizuno, Tomoaki, Naoki Hisamoto, Takashi Terada, Tae Kondo, Makoto Adachi, Eisuke Nishida, Dennis H. Kim, Frederick M. Ausubel, and Kunihiro Matsumoto. 2004. "The *Caenorhabditis Elegans* MAPK Phosphatase VHP-1 Mediates a Novel JNK-like Signaling Pathway in Stress Response." *The EMBO Journal* 23 (11): 2226–34.

Niu, Wei, Zhi John Lu, Mei Zhong, Mihail Sarov, John I. Murray, Cathleen M. Brdlik, Judith Janette, et al. 2011. "Diverse Transcription Factor Binding Features Revealed by Genome-Wide ChIP-Seq in *C. Elegans*." *Genome Research* 21 (2): 245–54.

Tang, Lanlan, William Dodd, and Keith Choe. 2016. "Isolation of a Hypomorphic *Skn-1* Allele That Does Not Require a Balancer for Maintenance." *G3 Genes|Genomes|Genetics* 6 (3): 551–58.
